# Using Biological Constraints to Improve Prediction in Precision Oncology

**DOI:** 10.1101/2021.05.25.445604

**Authors:** Mohamed Omar, Wikum Dinalankara, Lotte Mulder, Tendai Coady, Claudio Zanettini, Eddie Luidy Imada, Laurent Younes, Donald Geman, Luigi Marchionni

## Abstract

Many gene signatures have been developed by applying machine learning (ML) on *omics* profiles, however, their clinical utility is often hindered by limited interpretability and unstable performance in different datasets. Here, we show the importance of embedding prior biological knowledge in the decision rules yielded by ML approaches to build robust classifiers. We tested this by applying different ML algorithms on gene expression data to predict three difficult cancer phenotypes: bladder cancer progression to muscle invasive disease; response to neoadjuvant chemotherapy in triple-negative breast cancer, and prostate cancer metastatic progression. We developed two sets of classifiers: *mechanistic*, by restricting the training process to features capturing a specific biological mechanism; and *agnostic*, in which the training didn’t use any *a priori* biological information. Mechanistic models had a similar or better performance to their agnostic counterparts in the testing data, with enhanced stability, robustness, and interpretability. Our findings support the use of biological constraints to develop robust and interpretable gene signatures with high translational potential.

**Motivation:** *Omics*-based gene signatures often suffer from overfitting and reduced performance when tested on independent data. This usually results from the discrepancy between the high number of features compared to the much smaller number of samples used in the training process, which results in the machine learning algorithm perfectly fitting the training data with a subsequent deterioration in performance in independent cohorts. We introduce a mechanistic framework to mitigate overfitting and improve interpretability by constraining the training process to simple rank-based decision rules recapitulating relevant, cancer-related, biological mechanisms. Our approach aims at reducing the number of training variables to a pre-defined set of biologically important features in the form of gene pairs. The classification mechanism depends entirely on the relative ordering of these pairs, making it robust to data preprocessing techniques, improving the overall interpretability of the resulting models with significant translational implications. Most importantly, these pairs are configured in such a way that the decision rules resulting from the genes relative order embed and recapitulate specific biological mechanism, inherently enhancing the classifiers interpretability.

## Introduction

In oncology, machine learning (ML) algorithms are actively used to decipher gene expression data and identify predictive or prognostic gene signatures for specific cancer phenotypes like tumor progression or therapeutic response. Some of these signatures are currently being used in clinical settings to predict the prognosis and to guide further treatment (Cardoso et al., 2016; Knezevic et al., 2013). The process of gene signatures discovery and validation is hindered by relevant challenges (Mirza et al., 2019). The most striking one is the unstable performance of the discovered signatures when tested on different data than the ones used in their training. The main reason for this is the great discrepancy between the number of features or genes used for prediction (tens of thousands) and the number of observations or samples (tens to hundreds). In these settings, what can easily happen is that the ML model misinterprets “noise” as “signal” and ends up memorizing all the details in the training data which in turn cannot be generalized to other datasets, this is known as overfitting (Keogh and Mueen, 2010). There are several approaches to reduce overfitting and increase robustness, the most important of which is increasing the number of samples; however, this is not always feasible in biomedical research due to financial limitations or rarity of the studied disease phenotypes. Other options include using simple algorithms which are less susceptible to overfitting (Hand, 2006), using regularization with complex ones (Chicco, 2017; Neumaier, 1998), and reducing dimensionality by filtering out non-informative features or by using feature selection methods (Mahendran et al., 2020).

We hypothesize that embedding biological, mechanistic constraints in the decision rules during the training process will guide the ML algorithm to a set of features important for the phenotype being predicted, which in turn can reduce overfitting, and improve the performance, robustness, and interpretability of the resulting models. These constraints take the form of gene pairs which are derived from existing biological knowledge from the literature or pre-curated databases, and whose relative ordering determines the predicted class (Geman et al., 2004; Marchionni et al., 2013; Tan et al., 2005). Here, we test if this method can yield more interpretable gene signatures with a comparable performance to agnostic methods (*i.e.*, not based on prior biological knowledge) by using gene expression data to develop predictive classifiers in three distinct and hard prediction cases: 1. predicting the progression of non-muscle invasive bladder cancer (NMIBC) (stage T1) to muscle-invasive disease (MIBC) (stages T2-T4); 2. the response to neoadjuvant chemotherapy (NACT) in triple negative breast cancer (TNBC); and 3. prostate cancer (PCa) metastasis from primary tumor samples. In each of these three cases, we use four different ML algorithms: k-Top Scoring Pairs (k-TSPs), Support Vector Machine (SVM), Random Forest (RF), and Extreme Gradient Boosting (XGB).

To build mechanistic models, we restrict the training process to a specific biological mechanism relevant to the phenotype under study. For bladder cancer (BLCA) progression, we use feed-forward loops (FFLs) which consist of transcription factors (TFs) and microRNAs (miRNAs) target genes. TFs regulate the expression of their target genes through various mechanisms (Lambert et al., 2018), while miRNAs – a class of small, non-coding RNAs – play an important role in post-transcriptional gene regulation through the modulation of mRNA degradation (O’Brien et al., 2018). Current evidence shows that both TFs and miRNAs regulate the expression of common target genes and the expression of each other through feed-back (FBLs) and feed-forward loops (FFLs) (Friard et al., 2010; Hausser and Zavolan, 2014; Martinez et al., 2008; Re et al., 2009). Moreover, other studies have shown that the interaction between miRNAs targets and TFs is involved in the progression of several cancers including bladder cancer (Dong et al., 2017; Guo et al., 2013; Li et al., 2011; Liu et al., 2014; Mullany et al., 2018). Using the same principle for the TNBC case, we restrict the training process to mechanisms involving gene targets downstream to the Notch and MYC pathways, owing to the role these play in mediating cancer chemoresistance. Notch signaling pathway is involved in promoting cancer angiogenesis and epithelial-mesenchymal transition (EMT) (Abdullah and Chow, 2013). It also promotes chemoresistance in several cancers including breast cancer by inhibiting apoptosis and mediating cancer stem cells (CSC) self-renewal capacity (Ranganathan et al., 2011). Similarly, *MYC* promotes chemoresistance by mediating CSC self-renewal and proliferation (Wang et al., 2008; Zhang et al., 2019), and also by dysregulating the expression of some ATP-binding cassette (ABC) transporters necessary for cellular drug transport (Porro et al., 2010). Finally, since metastatic progression is mediated by several known biological processes, including loss of cell-cell adhesion and hypoxia in the tumor microenvironment (TME) (Bhandari et al., 2019; Oppenheimer, 2006), we designed a set of mechanistic pairs capturing such processes for predicting PCa metastasis.

In summary, here we embed prior knowledge of cancer biology directly into the algorithmic process to identify robust decision rules. We show that such mechanistic models, even with a relatively small number of features, have a similar or even superior performance and robustness, and an enhanced interpretability compared to agnostic models based on hundreds of genes solely selected based on statistical significance.

## Results

### Building mechanistic classifiers by embedding prior knowledge in the predictive decision rules

We hypothesized that integrating existing biological knowledge in the training process can yield robust and interpretable models the translational potential of which can surpass that of agnostic methods. For each classification task, we identified several biological processes related to the phenotype under study and used these to build corresponding biological mechanisms. To simplify the decision rules, we designed each mechanism as a list of gene pairs, each consisting of a gene associated with bad prognosis (*e.g.*, progression or chemo-resistance) and another associated with good prognosis. Specifically, for predicting BLCA progression, we built a mechanism based on feed-forward loops (FFLs) consisting of a TF which inhibits a downstream miRNA target gene (Figure S1). For the TNBC task, we based the mechanistic constraints on NOTCH and MYC signaling based on their involvement in mediating chemoresistance (Abdullah and Chow, 2013; Ranganathan et al., 2011; Wang et al., 2008; Zhang et al., 2019). Specifically, the mechanistic constraints were built by pairing the genes up-regulated with those down-regulated by NOTCH or MYC. Finally, for predicting PCa metastasis, we restricted the training process to gene pairs orchestrating cellular adhesion and O2 response.

We used such mechanistic pairs to train biologically-constrained, rank-based models (Geman et al., 2004; Tan et al., 2005), and then compared their performance to agnostic ones trained without biological constrains, starting from differentially-expressed genes or their pairwise combinations (Figure 1). Furthermore, we performed such a comparison using two distinct designs: training bootstrap and cross-study validation. In the bootstrap design, we proceeded as follows: we a) divided the data into training and testing sets; b) bootstrapped the training set 1000 times with a sample size equal to that of the training set (resampling with replacement); c) trained agnostic and mechanistic models on each training resample; and d) evaluated their performance on the untouched testing set. In the cross-study validation design, we used all but one study (*n-1)* for training, while the left-out study was used for testing, this process being repeated *n* times so that each study was used for testing.

**Figure 1.**
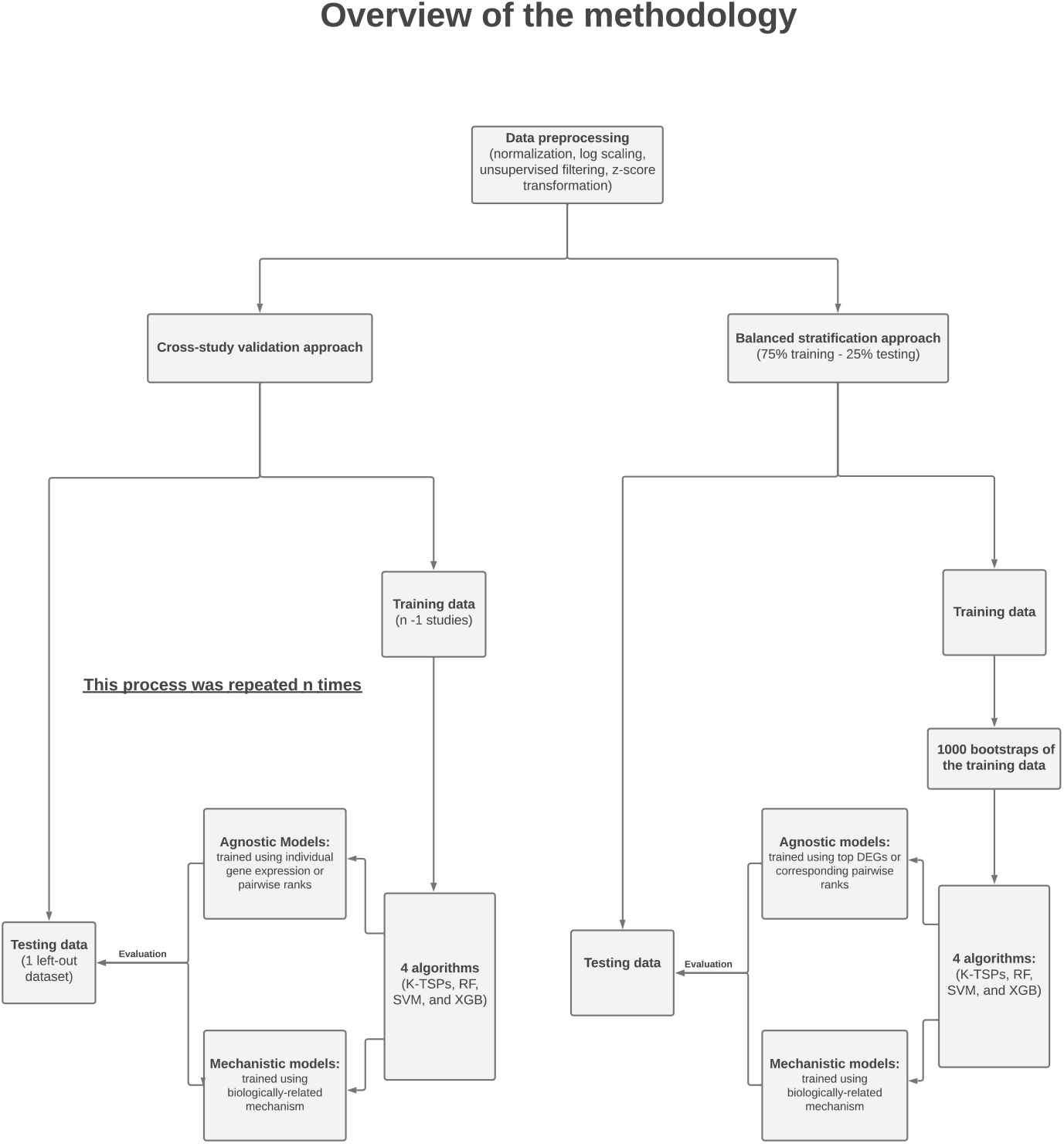
Building mechanistic classifiers by embedding prior knowledge in the classification decision rules. Three prediction cases were considered: predicting bladder cancer progression, the response to neoadjuvant chemotherapy in patients with triple-negative breast cancer, and prostate cancer metastatic progression. We adopted two different experimental designs: the training bootstrap and cross-study validation. In the bootstrap design, all datasets were pooled together after normalization and preprocessing, then split into training and testing sets. The training set was bootstrapped 1000 times and on each resample, we trained agnostic and mechanistic models then evaluated their performance on the testing set. In the cross-study validation, the analysis included *n* iterations where *n* corresponds to the number of studies. In each iteration, we used all, but one study for training the models and evaluated their performance on the left-out study. k-TSPs: k-top scoring pairs, RF: random forest, SVM: support vector machine, XGB: extreme gradient boosting, DEGs: differentially expressed genes.

### Mechanistic models based on FFLs outperform agnostic ones in predicting bladder cancer progression

In the BLCA task, we used gene expression profiles from 350 patients with NMIBC (stage T1) to train predictive models for cancer progression to MIBC (stage T2 or higher). We trained mechanistic classifiers using FFLs (37 unique mechanistic pairs) and compared their performance and robustness to agnostic models trained on the top differentially expressed genes, or the pairwise combinations of their ranks, without any prior biological consideration. Since the number of features used in the training process can be a major contributor to overfitting, we restricted the initialization of the agnostic models to the top 74 DEGs, matching the starting number of features used for training in the mechanistic case. Additionally, we also combined such top DEGs into 37 rank-based pairs to examine if this could improve the performance. In the bootstrap design, using the Area Under the ROC Curve (AUC) as evaluation metric, the agnostic and mechanistic k-TSPs models had a similar performance at predicting bladder cancer progression in the independent testing set (Figure 2). However, the mechanistic k-TSPs models were more parsimonious yielding on average five gene pairs compared to 16 pairs used by the agnostic ones. Importantly, the testing performance of the mechanistic models was more comparable with what was observed in the training. For the other three algorithms (RF, SVM, and XGB), the mechanistic models showed, on average, a higher testing performance than agnostic ones trained using the top DEGs (Figure 2). Interestingly, using pairwise comparisons derived from the top DEGs – instead of using their individual expression values – improved the performance of the agnostic models, slightly reducing the gap between training and testing. In secondary analyses, we also tested whether increasing the number of starting features in the agnostic case could improve their performance by training additional models using either the top 100, 200, and 500 DEGs or their pairwise comparisons (50, 100, and 250 pairs). Our results show that mechanistic classifiers still had a comparable or superior performance and robustness to agnostic ones, even when increasing the number of features (Figure S2). In addition, we also confirmed that mechanistic models held a clear advantage over agnostic ones trained using randomly selected genes (see Figure S3).

**Figure 2.**
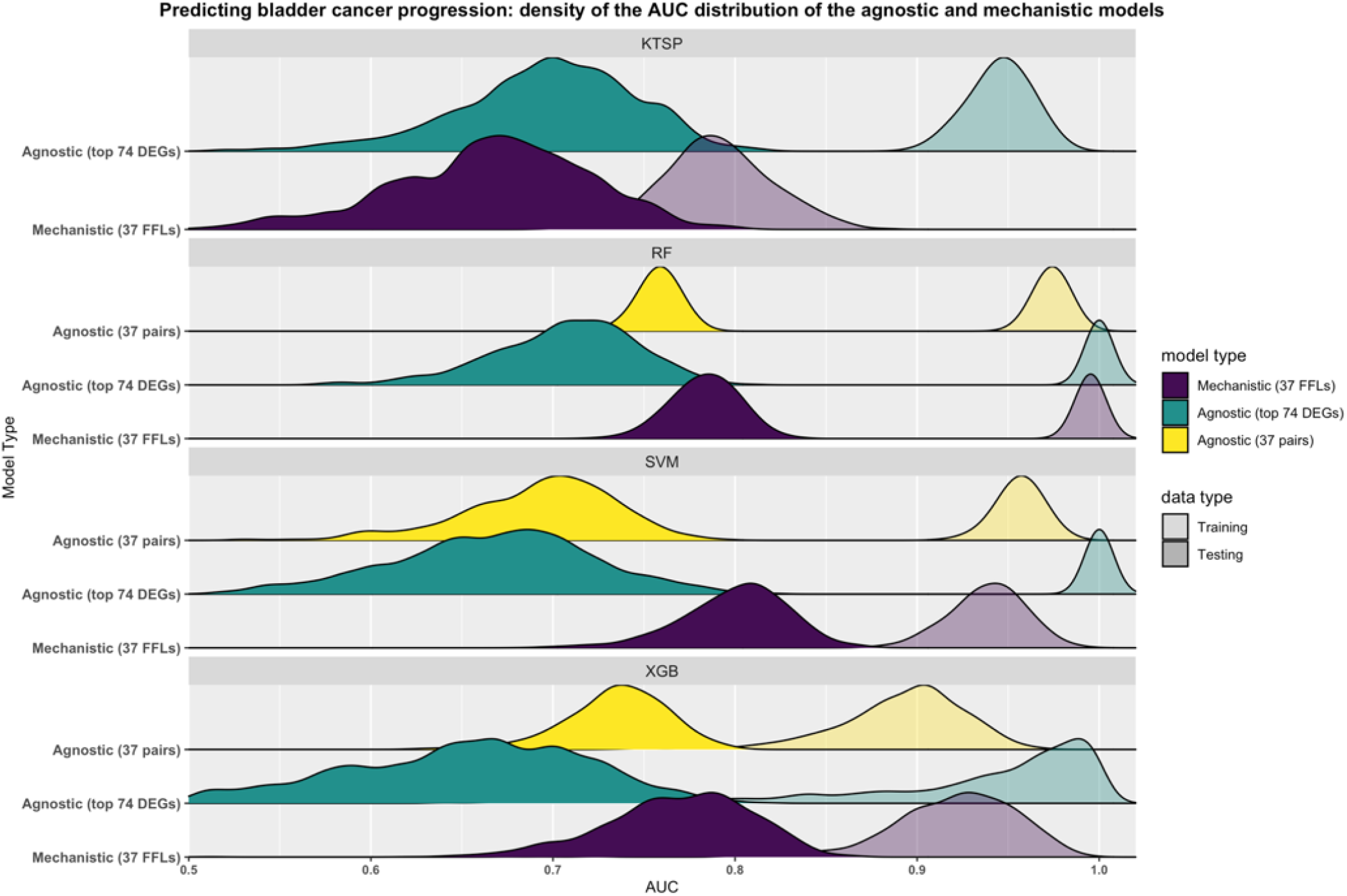
Mechanistic models based on FFLs outperform agnostic ones in predicting bladder cancer progression. The figure depicts the performance of the agnostic and mechanistic models as obtained using the described bootstrap design. Briefly, all models were trained on 1000 bootstraps of the training data (transparent colors), then evaluated on untouched testing data (solid colors) using the Area Under the ROC Curve (AUC) as performance metric. Mechanistic models were based on the feed-forward loops (37 pairs) (purple) and agnostic models were trained either using the top differentially expressed genes (74 genes) (green) or the corresponding pairwise comparisons (37 pairs) (yellow). Curves represent the smoothed density distributions of the AUC values, and each panel corresponds to one of the four algorithms used. KTSP: K-top scoring pairs; RF: random forest; SVM: support vector machine; XGB: extreme gradient boosting; FFLs: feed-forward loops; DEGs: differentially expressed genes.

To examine the consistency of gene signatures across the 1000 bootstraps, we ranked the gene pairs returned by the agnostic and mechanistic k-TSPs classifiers by their frequency. Interestingly, the mechanistic models tended to return more frequent pairs across the different training data resamples compared to the agnostic models, in which the selected pairs differed significantly with each training iteration (Figure 3A). For example, the most frequent gene pairs selected by the mechanistic k-TSPs models were ARHGEF11-RAD21 (n=564), CNOT3-RNF44 (n=422), CHD4-DFFA (n=380), HNRNPF-TP53 (n=324), and EPB41-GTF2B (n=280) (Figure 3B). On the other hand, the five most frequently selected pairs by the agnostic k-TSPs were MDC1-DECR1 (n=75), MDC1-USP31 (n=50), CPSF4-RPP40 (n=43), CHSY1-NIF3L1 (n=41), and SAC3D1-RPP40 (n=37) (Figure 3B). Each TSP gives a vote for a particular class (BLCA progression vs no-progression) if gene1 is more expressed than gene2. This means that the first gene in each pair can be interpreted as a gene associated with cancer progression, while the second would be expected to have an opposite role. With this notion in mind, we obtained – and then compared – the networks representing the frequency of each TSPs and that of each individual gene (across unique pairs), for both the agnostic and mechanistic k-TSPs approaches. The agnostic network was dominated by several genes associated with progression (mainly MDC1, RBP1, MMP11, CHD1L, and CDC25B) and no-progression (RPL12, RPP40, PSMB10, and PTGES2) (Figure 3C). In contrast, the mechanistic network was found to be more coherent, being dominated mainly by EZH2 serving as progression-associated gene (gene1) and RAD21, followed by TP53, serving as non-progression associated genes (gene2) (Figure 3D).

**Figure 3.**
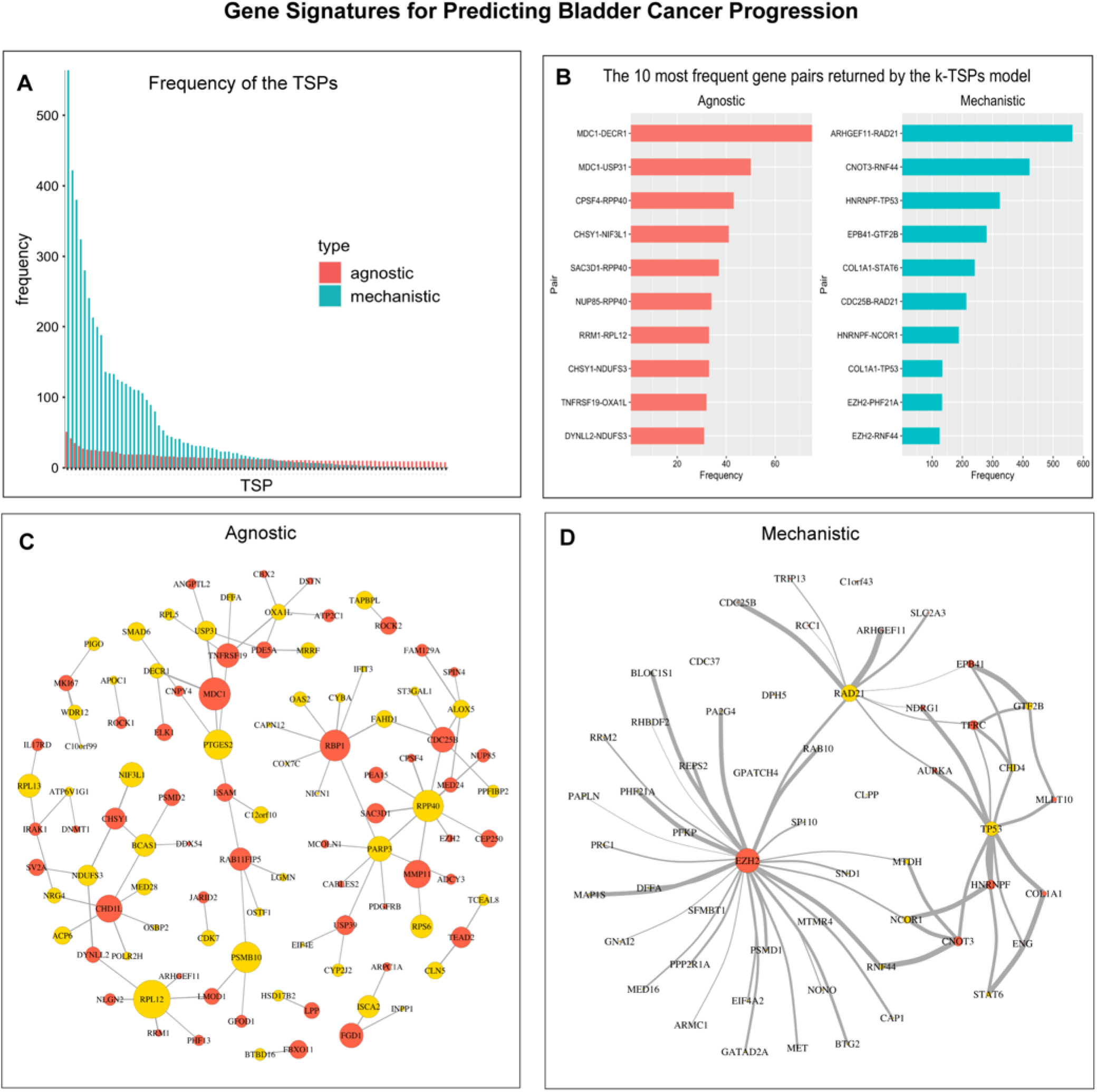
Mechanistic and agnostic k-TSPs signatures for predicting bladder cancer progression from non-muscle- to muscle-invasive stages. **A)** Bar plot showing the frequency of the agnostic (red) and mechanistic (blue) scoring pairs across the different bootstraps. The bar plot includes all the mechanistic pairs (n=93) and the most frequent agnostic pairs (n=93), both sorted by their frequency across the different bootstraps. **B)** The top 10 most frequent agnostic (red) and mechanistic (blue) pairs sort by their frequency. **C-D)** Networks of the top 93 agnostic (C) and mechanistic (D) top-scoring pairs. Each pair consists of a gene voting for BLCA progression (red) and no progression (yellow). The vertex size corresponds to 2*log2 of the individual gene frequency across unique pairs while the edge thickness corresponds to the log2 of the top scoring pair frequency.

To examine the functional states of genes associated with BLCA progression compared to those associated with no-progression, we performed gene set enrichment analysis (GSEA) on all the genes positioned as either gene1 (*i.e., the “bad”* genes) or gene 2 (*i.e., the “good”* genes) in the TSPs, separately for mechanistic and agnostic classifiers. Interestingly, genes in the mechanistic signatures (55 unique genes) were significantly enriched in 62 biological processes, many of which are related to invasion and progression including EMT, cell proliferation, and cell cycle transition (Table S1). On the other hand, genes in the agnostic signatures were significantly enriched in only one biological process despite their larger number (74 unique genes) (Table S1).

Finally, in the cross-study validation design, we found that both the mechanistic and agnostic k-TSPs models had a similar average AUC in the testing data, while the mechanistic one outperformed its agnostic counterpart on multiple metric (balanced accuracy, sensitivity, and MCC (Chicco et al., 2021), see Table S2). In agreement with the bootstrap results, the testing performance of the mechanistic k-TSPs was also highly comparable with that of the training, suggesting improved generalizability. Similar results were also seen with the other three ML algorithms (RF, SVM, and XGB, see Table S2).

### NOTCH-MYC-based models outperform their agnostic counterparts in predicting response to neoadjuvant chemotherapy in TNBC

For predicting the response to NACT in patients with TNBC, we used the pre-treatment gene expression profiles of 369 patients with TNBC. Mechanistic models were trained using the NOTCH-MYC mechanism (241 unique pairs), while agnostic models were trained using the top 500 DEGs, or their corresponding pairwise-ranks (250 pairs). Performance was compared with the same two designs described for the bladder cancer case.

In the bootstrap approach, our results show that the mechanistic k-TSPs models have similar testing performance to the agnostic ones, however, the mechanistic RF, SVM, and XGB still slightly outperformed their agnostic counterparts (Figure 4). Furthermore, changing the number of training features only improved the testing performance of the agnostic models based on pairwise-ranks, while still falling short of the mechanistic ones (Figure S4). Finally, mechanistic models clearly outperformed those trained on random genes (Figure S5).

**Figure 4.**
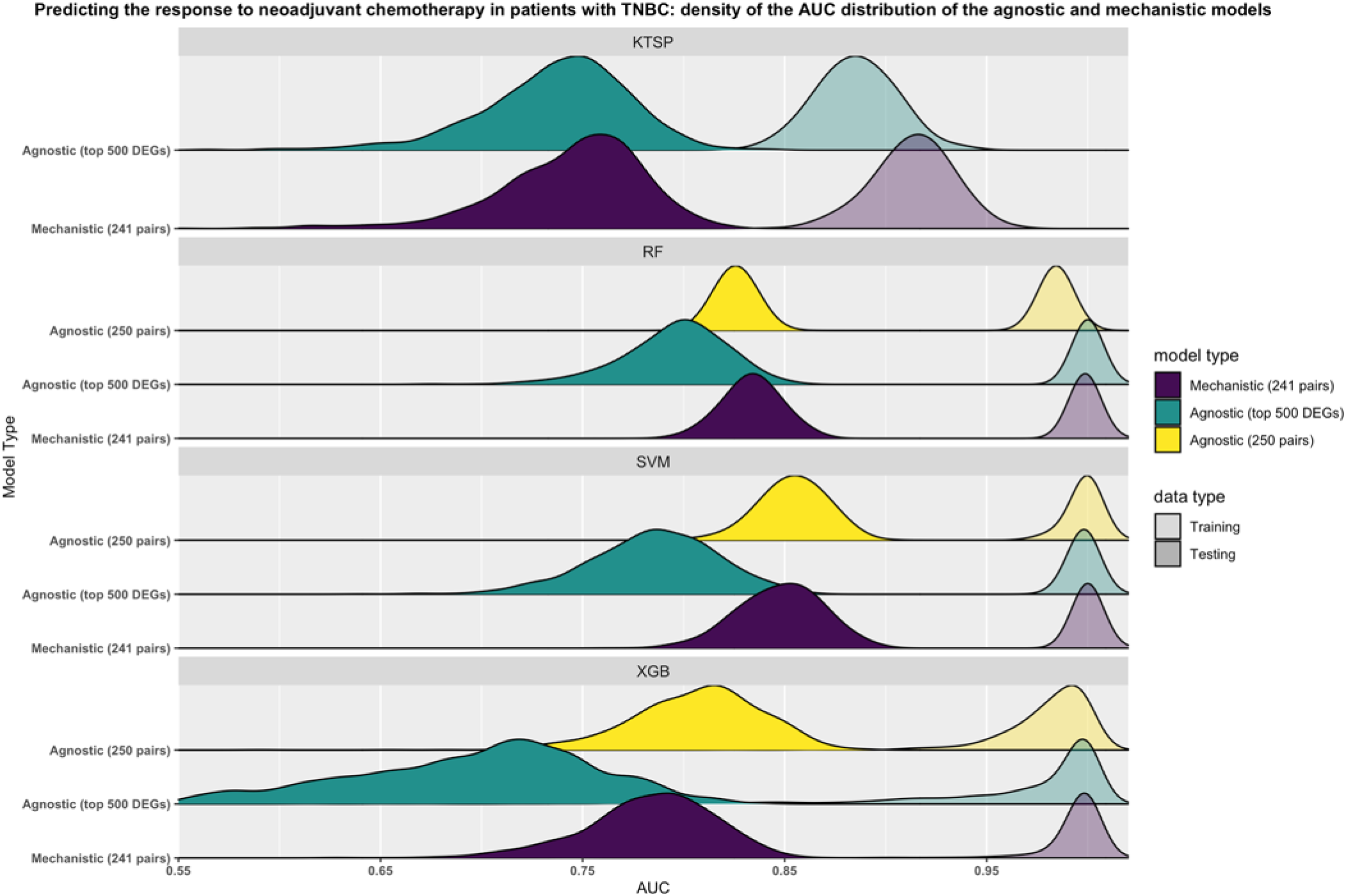
NOTCH-MYC-based models outperform their agnostic counterparts in predicting response to neoadjuvant chemotherapy in patients with triple-negative breast cancer. Models were trained on 1000 bootstraps of the training data (transparent colors) and evaluated on the untouched testing data (solid colors) using the Area Under the ROC curve (AUC) as metric. Mechanistic models were based on the NOTCH-MYC mechanism (241 pairs) (purple) while agnostic models were trained either using the top differentially expressed genes (500 genes) (green) or the corresponding pairwise comparisons (250 pairs) (yellow). Shown are the smoothed density distributions of the AUC values with each panel corresponding to one of the four algorithms used. KTSP: K-top scoring pairs; RF: random forest; SVM: support vector machine; XGB: extreme gradient boosting; DEGs: differentially expressed genes; TNBC: triple-negative breast cancer.

Ranking out pairs by their frequency across the different iterations showed that a similar and more robust set of mechanistic pairs get selected much more often than agnostic ones (Figure 5A). For instance, the five most frequently returned pairs by the mechanistic k-TSPs models included DDIT3-DDX18 (n=531), TSC2-PLK4 (n=499), COL5A1-ITGA6 (n=442), GARS-PDCD10 (n=385), and NDE1-EZR (n=347) (Figure 5B). However, the five most frequent pairs returned by the agnostic k-TSPs models included SLC43A1-ABT1 (n=204), ITGA5-EPHB3 (n=119), METRN-MCM5 (n=116), PARM1-MAPK9 (n=97), and SLC22A5-DCAF7 (n=96) (Figure 5B). Here, the decision rules follow the same pattern discussed in the BLCA case, with each pair voting for either residual disease (RD) or pathological complete response (pCR) based on the expression of the two genes. In agnostic models, the most frequent gene associated with RD (gene1 in the TSP) was MAST3 while RPL39L was the most frequent gene associated with pCR (gene2 in the TSP) (Figure 5C). For the mechanistic models, the oncogene CCND1 was the most frequent gene overexpressed in samples from patients with RD (gene1 in the TSP) while the WNT antagonist SFRP1 (Veeck et al., 2006) was the most frequently overexpressed in samples from patients who had pCR (gene2 in the TSP) (Figure 5D).

**Figure 5.**
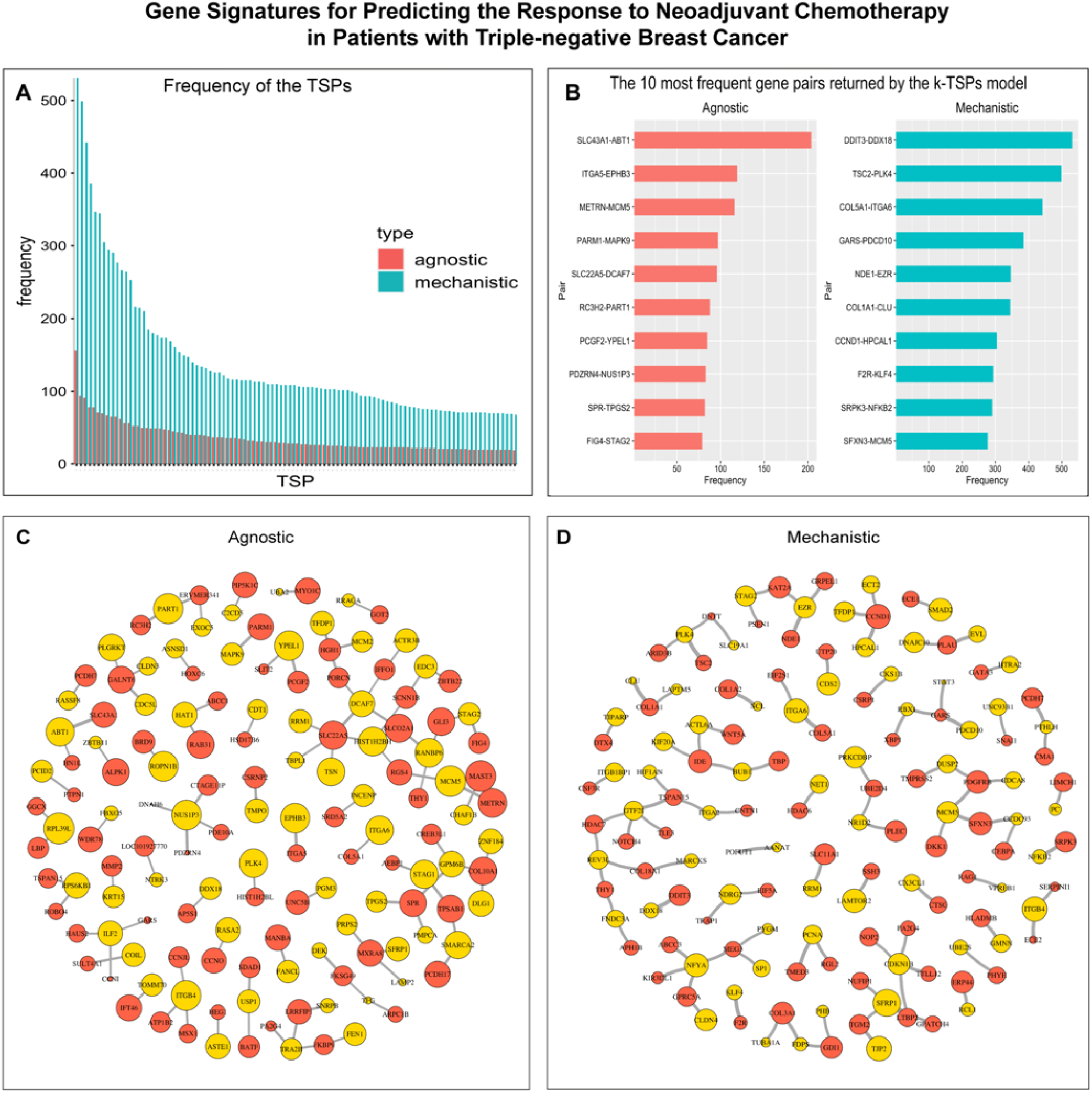
Mechanistic and agnostic k-TSPs signatures for predicting the response to neoadjuvant chemotherapy in patients with triple-negative breast cancer. **A)** Bar plot showing the frequency of the top 100 agnostic (red) and mechanistic (blue) scoring pairs across the different bootstraps. **B)** The top 10 most frequent agnostic (red) and mechanistic (blue) pairs sort by their frequency. **C-D)** Networks of the top 100 agnostic (C) and mechanistic (D) top-scoring pairs. Each pair consists of a gene voting for RD (red) and pCR (yellow). The vertex size corresponds to 2*log2 of the individual gene frequency across unique pairs while the edge thickness corresponds to the log2 of the top scoring pair frequency.

The genes from the mechanistic classifiers (656 unique genes) were significantly enriched in 780 different pathways and processes including those used to build the priori mechanism (NOTCH and MYC signaling) (Table S3). They were also enriched in other cancer-related pathways like regulation of apoptosis, beta-catenin-TCF complex assembly, TGF-B signaling, T-cells activation and differentiation. While genes from agnostic classifiers were larger in number (1335 unique genes), they were significantly enriched in only 49 gene sets without a strong association with cancer biology. Altogether, these results reflect the signatures selected by the k-TSPs models are consistent and more associated with the biological processes underlying chemotherapeutic resistance compared to those returned by agnostic models (Table S3).

Lastly, also in the cross-study validation case, we did not observe significant differences in performance between the model types, however, the mechanistic classifiers showed slightly more consistency in performance metrics like the AUC between training and testing, especially using the k-TSPs and XGB algorithms (Table S4). While using biological constraints did not offer a clear advantage in terms of performance, it significantly enhanced interpretability, and reduced the computational costs by limiting the training to a few hundred features instead several thousand used for the agnostic models.

### Mechanistic models based on cellular adhesion and oxygen response have a similar performance to their agnostic counterparts in predicting prostate cancer metastatic progression

For predicting metastatic progression in prostate cancer, we used seven gene expression datasets comprising 1239 primary tumor samples including 399 with metastatic events.

In the bootstrap approach, mechanistic models using 50 pairs had a similar performance to their agnostic counterparts and incurred in less overfitting irrespective to the number of training features (Figure 6). Additionally, these mechanistic classifiers still maintained their performance even when more features were used for training the agnostic models (Figure S6), or when random genes were used in the training process (Figure S7).

**Figure 6.**
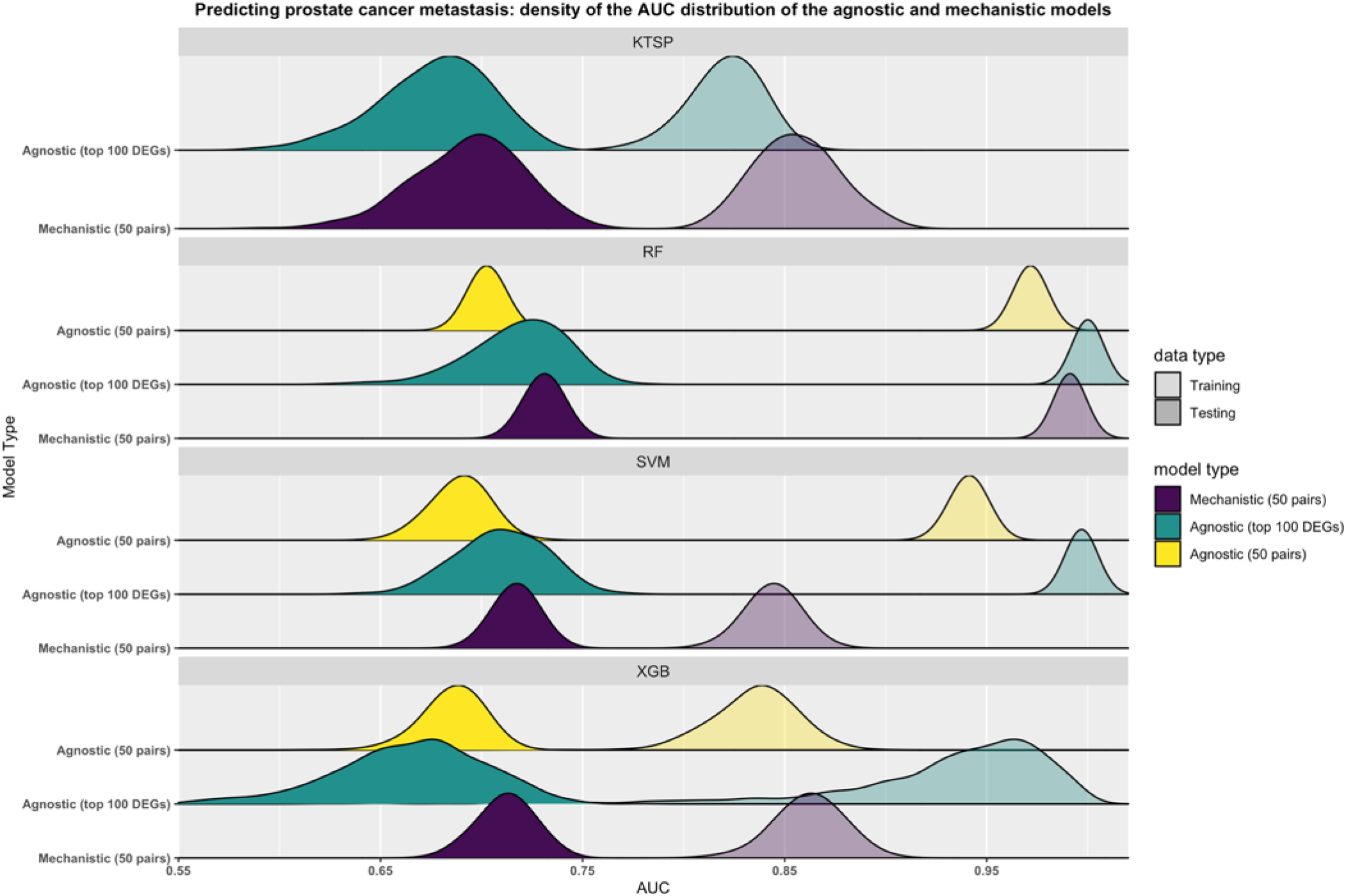
Mechanistic models based on cellular adhesion and oxygen response have similar performance to their agnostic counterparts in predicting prostate cancer metastatic progression. The figure depicts the results from the bootstrap design in which the training set (transparent colors) was resampled 1000 times. On each resample, models were trained to predict metastatic progression in prostate cancer and their performance was evaluated on the untouched testing set (dark colors) using the Area Under the ROC Curve (AUC) as evaluation metric. Mechanistic models were based on the cellular adhesion and O2 response mechanism (50 pairs) (purple) while agnostic models were trained using either the top differentially expressed genes (100 genes) (green) or the corresponding pairwise comparisons (50 pairs) (yellow). Shown are the smoothed density distributions of the AUC values and each panel corresponds to one of the four algorithms used. KTSP: K-top scoring pairs; RF: random forest; SVM: support vector machine; XGB: extreme gradient boosting; DEGs: differentially expressed genes.

Mechanistic TSPs showed high frequency (Figure 7A) with the five most frequent pairs including S100A10-PTN (n=540), CD74-SATB1 (n=455), CBX3-AZGP1 (n=414), CXCR4-PCDH18 (n=237), and STAT1-DPP4 (n=214) (Figure 7B). Agnostic models on the other hand frequently returned CAMK2N1-CDC42EPS (n=389), CXCR4-LPAR3 (n=230), ENO1-CDC42EPS (n=217), RFTN1-DPT (n=202), and GNPTAB-CTBS (n=190) (Figure 7B-C). Interestingly, in the mechanistic TSPs, genes related to PCa progression and metastases like THBS2 (Chen et al., 2017), NRP1(Tse et al., 2017), and WNT5A (Dai et al., 2008) were frequently represented as metastases-voting genes (gene1 in the TSPs) (Figure 7D).

**Figure 7.**
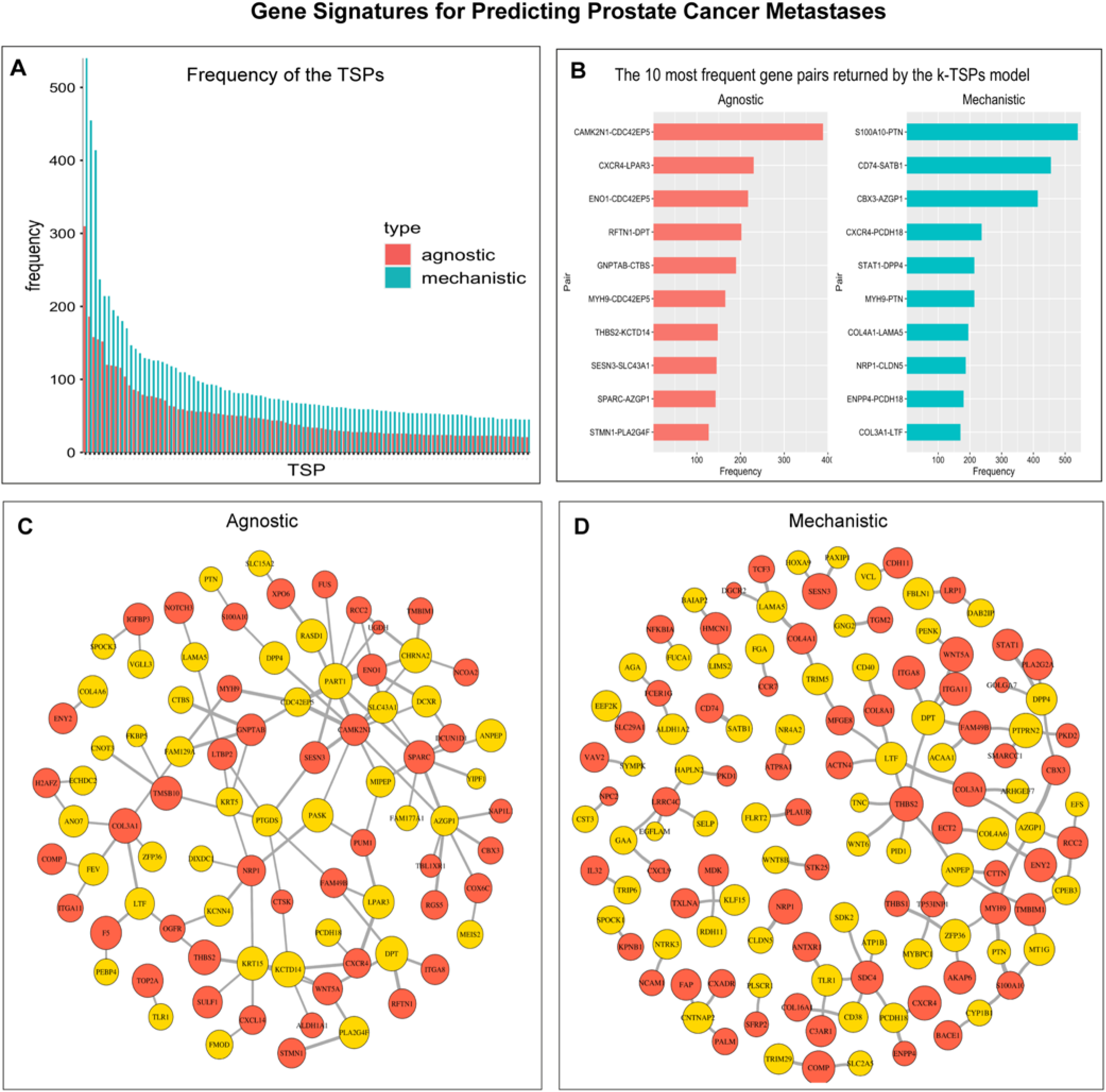
Mechanistic and agnostic k-TSPs signatures for predicting prostate cancer metastases. **A)** Bar plot showing the frequency of the top 100 agnostic (red) and mechanistic (blue) scoring pairs across the different bootstraps. **B)** The top 10 most frequent agnostic (red) and mechanistic (blue) pairs sort by their frequency. **C-D)** Networks of the top 100 agnostic (C) and mechanistic (D) top-scoring pairs. Each pair consists of a gene voting for metastasis (red) and no-metastasis (yellow). The vertex size corresponds to the 2*log2 of the individual gene frequency across unique pairs while the edge thickness corresponds to the log2 of the top scoring pair frequency.

Gene set Enrichment analysis showed that genes derived from the mechanistic signatures (906 unique genes) were significantly enriched in 1920 gene sets, many of which are associated with cell migration, motility, adhesion, and proliferation (Table S5). They were also enriched in other important pathways involved in PCa progression and metastases including regulation of MAPK and ERK cascades, NF-kappaB, STAT, TGF-beta, and RAS signaling pathways. Similarly, genes from the agnostic models (1622 unique genes) were enriched in 96 gene sets, some of which included regulation of cell migration and motility and PCa related pathways like WNT signaling (Table S5).

Finally, these results were also confirmed in the cross-study validation analysis, in which the mechanistic models had similar performance compared to the agnostic ones, but provided superior interpretability and improved computational efficiency, due to reduced number of features (tens versus thousands) used in the training process (Table S6).

### The right mechanism for the right task: mechanistic constraints should be related to the phenotype under consideration

We have shown that using biological constraints in the training process can improve the performance of the resulting gene signatures, however, we hypothesized that this is contingent on using a mechanism related to the phenotype under-study. To investigate this, we assessed the performance of several different mechanisms in each of the three classification tasks we considered. This analysis included the three afore-mentioned cancer-related mechanisms (FFLs, NOTCH-MYC signaling, and cellular adhesion and O2 response), together with three other, non-cancer related ones, constructed from molecular profiles involved in: a) Alzheimer disease (266 pairs); b) diabetes (6776 pairs); and c) viral infection (6384 pairs).

For each cancer phenotype, models trained on the relevant mechanism of choice showed the best performance (Table 1). Specifically, classifiers trained using the FFLs had higher testing AUCs for predicting bladder cancer progression compared to those based on the other mechanisms. Similar results were also obtained for NACT response prediction in TNBC using NOTCH-MYC signaling, and for prostate cancer metastatic progression using cellular adhesion and O2 response.

**Table 1.**
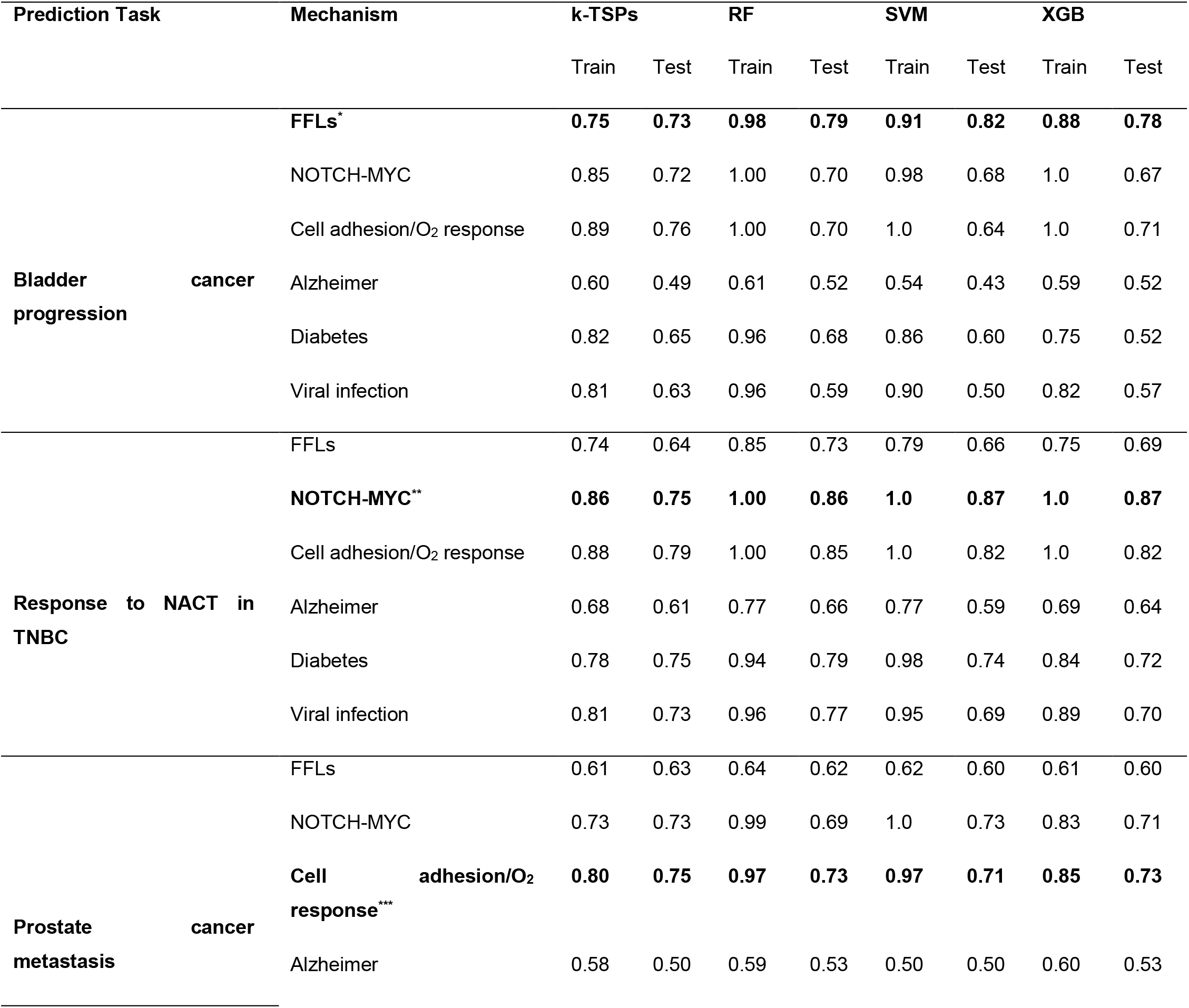
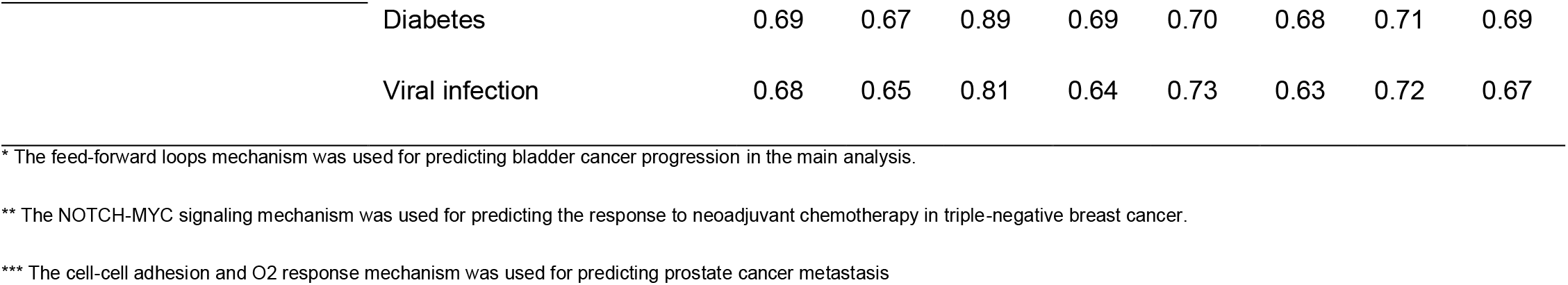
The performance of different mechanisms at predicting each of the three cancer phenotypes assessed by the Area under the ROC curve (AUC). Three important cancer phenotypes were considered for prediction: bladder cancer progression from non-muscle invasive to muscle-invasive stages, response to neoadjuvant chemotherapy (NACT) in patients with triple-negative breast cancer (TNBC), and metastatic progression in prostate cancer. For each phenotype, we built a priori mechanism capturing the underlying biology (bold text) and used it for prediction in the main analysis. Feed-Forward Loops (FFLs) were designed for predicting bladder cancer progression while the NOTCH-MYC signaling and cellular adhesion and O2 response mechanisms were developed for predicting the response to NACT in TNBC and prostate cancer metastatic progression, respectively. We also collected three cancer unrelated mechanisms and used them as negative controls. These include the Alzheimer, diabetes, and viral infection mechanisms. The performance of the different cancer-related and unrelated mechanisms at predicting each of the three cancer phenotypes was assessed using the Area Under the ROC Curve (AUC). FFLs: feed-forward loops, NACT: neoadjuvant chemotherapy, TNBC: triple-negative breast cancer.

Notably, the three cancer related mechanisms maintained a superior performance over the non-cancer related ones in each of the three prediction tasks. For example, while the FFLs-based classifiers had the best performance at predicting bladder cancer progression, the NOTCH-MYC signaling and cellular adhesion/O2 response mechanisms also achieved a good testing performance. On the other hand, the Alzheimer, diabetes, and viral infection mechanisms performed poorly in the three classification cases (Table 1), highlighting the fact that choosing the right mechanism for the prediction task is essential for achieving an optimal performance.

## Discussion

Overfitting and lack of generalization remain among the most difficult problems in machine learning especially in transcriptomics owing to the very large number of features and the much smaller number of samples (Keogh and Mueen, 2010; Mirza et al., 2019). The inconsistency of performance of genomic predictive models is one of the reasons why their clinical usage has not been widely implemented, and it represents a major obstacle towards personalizing health care. While some approaches like increasing the sample size may help improve the performance and stability of such models, these may not always be feasible in medical research, owing to financial limitations or the unavailability of samples in cases of rare cancer phenotypes.

Some studies have shown that using prior knowledge can help to choose the input data or the correct algorithm (Libbrecht and Noble, 2015; Yip et al., 2012). In this study, we employed a similar concept to train robust and interpretable predictive models by adding biological constraints to the decision rules used for classification. We examined this approach in three clinically important classification cases: predicting bladder cancer progression from non-muscle invasive to muscle invasive stages, predicting the response to neoadjuvant chemotherapy in TNBC, and predicting prostate cancer progression to metastasis disease. In each setting, we employed multiple algorithms and compared the performance of mechanistically constrained models to agnostic ones. In the bladder cancer case, we used FFLs based on the evidence supporting their involvement in cancer progression and invasion (Dong et al., 2017; Guo et al., 2013, p.; Li et al., 2011; Liu et al., 2014; Mullany et al., 2018). In the TNBC case, we focused on NOTCH and c-MYC targets based on their role in mediating cancer stem cells self-renewal and chemo-resistance (Ranganathan et al., 2011; Wang et al., 2008; Zhang et al., 2019). Finally, for predicting prostate cancer metastasis, we used gene pairs capturing cell-cell adhesion and the cellular response to O2 (Bhandari et al., 2019; Oppenheimer, 2006). Since each of these priori mechanisms is already known to be associated with the corresponding phenotype, the features constituting the mechanism are expected to be of high quality for the prediction task undertaken. In this case, the ML algorithm is essentially used to extract the smallest number of features that can serve as a gene signature and has more potential to generalize to other datasets. It is true that agnostic models can also extract a number of features with a similar performance if not better. However, in that case, the search process would be applied on all genes and has more risk to overfit to the training data and select non-informative features that can’t be generalized. When evaluated on the testing data, the mechanistic models yielded a similar or superior performance than the agnostic ones. Moreover, especially in the bladder cancer case, and for the K-TSP algorithm, mechanistic models demonstrated better generalization from training to testing, even in situations where there was a small number of samples available for training. Furthermore, these results were not dependent on any specification or characteristics pertaining to the training data, as shown by results from both the bootstrap and the cross-validation analyses.

It is important to note that the mechanism used in the training should be biologically related to the phenotype being predicted. We demonstrated this by assessing the performance of different mechanisms in each classification case and noticed that the mechanisms we designed based on prior knowledge had the best performance at predicting their corresponding phenotype. Interestingly, the three cancer related mechanisms we have considered achieved optimal performance in each cancer phenotype, while the non-cancer ones performed poorly, which further support our rationale for using prior related biological knowledge to develop robust classifiers.

Overall, these results show that models using a small number of biologically important features can have a similar or better performance metrics compared to those using hundreds or thousands of genes. Furthermore, mechanistic rank-based decision rules greatly enhance the interpretability of the resulting predictive models, which is crucial to their clinical adaptation. Specifically, the priory mechanism is designed beforehand by pairing genes from pathways positively and negatively associated with the phenotype of interest. In this case, the mechanism consists of multiple gene pairs belonging to opposing pathways or biological processes. Subsequently, we use the k-top scoring pairs (k-TSPs) algorithm which is rank based and serves by selecting the gene pairs whose expression consistently switch between the two classes of interest. For example, the NOTCH mechanism consists of hundreds of gene pairs up- and down-regulated by NOTCH signaling and we use this mechanism to design a classifier to predict the response to chemotherapy in breast cancer. The resulting classifier consists of a number of gene pairs with each consisting of a gene up- and another down-regulated by NOTCH signaling. Each of these pairs votes for a particular class (sensitive versus resistant) based on the order of expression of the two genes and the final prediction of a patient/sample is determined by the majority of votes. As such, the classifier is interpretable because its decision rules and the function of its genes are known beforehand. Also, such models are more robust to preprocessing techniques, and can be more easily implemented using other technologies already being used for clinical applications. For example, we have shown that such mechanistic signatures derived from microarrays or RNA-Seq studies can be implemented using technologies like RT-PCR, which further increases their translational value (Ghantous et al., 2022).

Together with their good performance, the mechanistic signatures captured the underlying biology of the associated cancer phenotype as expected. Since we used TF-miRNA targets as a priori mechanism to train ML models capable of predicting bladder cancer progression, the resulting signatures were always gene pairs with the 1^st^ gene being a TF while the second is a miRNA target gene mirroring the priori mechanism. Even with different training data resampling, these signatures were often consistent indicating their predictive value rather than perfectly fitting the training data. For instance, ARHGEF11-RAD21 was present in more than 50% of the mechanistic BLCA progression signatures voting for progression if ARHGEF11 is more expressed relative to RAD21. While the role ARHGEF11 in bladder cancer is not well-defined, it was found to be associated with the invasiveness of progression of glioblastoma (Ding et al., 2018) and hepatocellular carcinoma (Du et al., 2020). Across all mechanistic pairs, the transcriptional repressor EZH2 was prominent as a progression voting gene being present in 41 unique TSPs in which its expression relative to the second gene determines the vote given by the TSP (progression if EZH2 is more expressed than gene2). This is consistent with existing evidence linking EZH2 to progression and poor prognosis in several cancers including BLCA (Wang et al., 2012; Zhou et al., 2018). Similarly, predicting the response to NACT in patients with TNBC relied on a priori mechanism of gene pairs regulated by NOTCH and MYC signaling. Expectedly, the resulting signatures consisted of a small subset of gene pairs up- and down— regulated by either NOTCH or MYC, and among these, genes down-regulated by either TF (*e.g.*, SFRP1, ITGB4, and ITGA6) were predominantly associated with pCR. The same pattern was seen in predicting PCa metastases in which the mechanistic signatures frequently included genes known to be involved in mediating EMT and distant metastases e.g., THBS2 (Chen et al., 2017) and WNT5A (Dai et al., 2008). Gene set enrichment analyses showed that genes from the mechanistic signatures were significantly enriched in many more coherent biological processes and pathways compared to the genes from their agnostic counterpart, despite the latter being in much larger numbers. Moreover, In the three classification tasks, genes from the mechanistic classifiers were enriched in the pathways used to build the priori mechanism (as to be expected), together with other pathways associated with the phenotype under study. Altogether, these results show that mechanistic signatures tend to capture important cancer biology related to their corresponding phenotypes.

It is important to note that our study has some inherent limitations. First, the biological mechanisms we used take the form of contrasting gene pairs, but this pairwise relationships may not completely capture the complexity of the underlying biology compared to other formulations like gene networks. However, this lack of sophistication in the design of the biological mechanism was deemed necessary for the interpretability of the resulting predictive models. Moreover, such pairwise comparisons, which are used by the k-TSPs algorithm, can also be used as input to other more complex approaches (e.g., SVM, RF, and XGB), with the advantage of increasing the level of interpretability of the resulting models. Finally, while our study was focused on cancer, the same conceptual framework can be also applied to other diseases, provided there is available prior knowledge about the underlying pathophysiological mechanisms.

Despite these limitations, our work supports the adoption of mechanistically constrained decision rules for the development of robust prognostic and predictive models. Their high performance and intrinsic interpretability will promote a wider integration into clinical practice, bringing routine personalized medicine one step closer to reality.

## ACKNOWLEDGMENTS

We thank Dr. Soren Vang (Department of Molecular Medicine, Aarhus University Hospital, Denmark) for providing the raw RNA-Seq counts of the E-MTAB-4321 dataset.

## Author contributions

LM and DG conceived the research question. MO, LM, TC, CZ, ELI, and WD collected the datasets and gene sets. MO performed the analysis and wrote the manuscript. LM, DG, and LY supervised the analysis and the manuscript writing. All authors read and approved the final version of the manuscript.

## STAR Methods

### Resource availability

#### Lead contact

Further information and requests for resources and reagents should be directed to the lead contact, Dr. Luigi Marchionni.

#### Material Availability

This study did not generate new materials.

#### Data and Code Availability

This paper analyzes existing, publicly available data. The accession numbers for the datasets are listed in the key resources table. All original code together with our curated biological mechanisms have been deposited at https://github.com/MohamedOmar2020/Biological_Constraints and are publicly available.

### Method details

#### Data collection

##### Bladder cancer

We used both the NCBI Gene Expression Omnibus (GEO) (Barrett et al., 2007) and ArrayExpress (Kolesnikov et al., 2015) to identify gene expression datasets containing primary tumor samples from non-muscle invasive bladder cancer (NMIBC). We refined the initial results to keep only the datasets with information about the progression status (progression to MIBC versus no progression). Five datasets met our inclusion criteria, four of which are microarray-based (GSE57813 (van der Heijden et al., 2016), GSE13507 (Kim et al., 2010), GSE32894 (Sjodahl et al., 2012) and pmid15930337 (Dyrskjøt et al., 2005)) and the fifth is RNA-Seq based dataset (E-MTAB-4321 (Hedegaard et al., 2016)).

##### Breast cancer

We identified seven datasets with pre-treatment gene expression profiles from patients with breast cancer who received neoadjuvant chemotherapy: GSE25055 (Hatzis et al., 2011), GSE25065, GSE140494 (Edlund et al., 2021), GSE103668 (Birkbak et al., 2018), GSE20194 (Popovici et al., 2010; Shi et al., 2010), GSE20271 (Shen et al., 2012; Tabchy et al., 2010), and GSE32646 (Miyake et al., 2012).

##### Prostate cancer

For predicting PCa metastatic progression, we identified seven gene expression datasets containing primary tumor samples from PCa patients: GSE116918 (Jain et al., 2018), GSE55935 (Ragnum et al., 2015), GSE51066 (Ross et al., 2014), GSE46691 (Erho et al., 2013), GSE41408 (Boormans et al., 2013), JHU natural history cohort (Ross et al., 2016), and GSE70769 (Ross-Adams et al., 2015).

#### Data preprocessing

##### Bladder cancer

In each dataset, we removed non-invasive papillary carcinoma (Ta) and carcinoma in situ (Tis) samples and kept only T1 lesions with information about the progression status. To remove uninformative features in the microarray datasets, we kept genes with raw intensity greater than 100 in at least 50% of the samples. Similarly, in E-MTAB-4321 (RNA-Seq) we kept genes with more than one count per million (CPM) in at least 50% of the samples. The four microarray datasets were normalized and log2-scaled upon retrieval from GEO. For E-MTAB-4321, the read counts were normalized using trimmed mean of M-values (TMM) and transformed to log2-counts per million (log-CPM). Next, we performed Z-score transformation (by gene) of each normalized dataset separately to ensure that the datasets from both technologies (microarrays and RNA-Seq) are on a similar scale.

##### Breast cancer

The seven breast cancer datasets originally included 1013 samples which we filtered to keep only samples in which ER, PR, and HER2 were all negative by immunohistochemistry (IHC) and with available information about the response to NACT whether pathological complete response (pCR) or residual disease (RD). This reduced the number of samples used in downstream analysis to 369 TNBC samples. All datasets were normalized and log2-scaled when retrieved from GEO, except for GSE20271 in which the expression values were not logged and so was log2-transformed. Finally, we mapped each probe ID to the corresponding gene symbol and filtered the expression matrices to the gene symbols in common (13299 genes).

##### Prostate cancer

In each of the seven PCa datasets, we kept primary tumor samples with information about metastatic events resulting in 1239 primary tumor samples eligible for downstream analysis. Normalization and preprocessing were performed as described above followed by z-score transformation for each dataset separately. Finally, probe IDs were mapped to their corresponding gene symbols and expression matrices were restricted to the genes in common.

##### Platform harmonization

For the prediction tasks in bladder and prostate cancers, which included data obtained across different platforms and technologies (microarrays and RNAseq), we harmonized the final datasets by identifying a subset of cross-study reproducible genes using integrative correlation coefficient (ICC) (Cope et al., 2014; Parmigiani et al., 2004) keeping only genes whose ICC was greater than 0.15 or the 33rd percentile. In summary, the ICC is computed by calculating the Pearson correlation coefficient of the expression values of each pair of genes within and across studies (correlation of correlation). Although the integrative correlation analysis was performed on all data before division into training and testing, this does not violate the validation process since this method only uses the expression data and does not consider the phenotype information. Using the ICC threshold mentioned above, 3109 and 4055 genes were identified and used in the bladder and prostate cases, respectively.

#### Mechanistic pairs assembly

##### Feedforward loops

The TF-miRNA mediated gene regulatory loops that we are interested in are the coherent feed-forward loops in which a TF (e.g., MYC) inhibits a target gene (e.g., CD164) directly and indirectly via activation of a hub miRNA (e.g., has-miR-346). The TF and target gene have an inverse relationship; over-expression of the TF results in down-regulation of the target gene and vice versa (Figure S1). This inverse relationship makes these pairs suitable for classification. To construct these loops, three different interaction types must be obtained: the interaction between the TF and target gene (TF-target), the interaction between the TF and miRNA (TF-miRNA), and the interaction between miRNA and target gene (miRNA-target). The TF-target interactions were obtained from Harmonizome (Rouillard et al., 2016) using the following databases: ENCODE, ESCAPE, CHEA, JASPAR, MotifMap and TRANSFAC. The TF-miRNA interactions were obtained from the same databases as above together with TransmiR v2.0 database(Tong et al., 2019). Finally, the miRNA-target interactions were obtained from TargetScan (Agarwal et al., 2015), miRTarBase (Chou et al., 2021) and miRWalk (Sticht et al., 2018).

The loops were constructed by merging the three different interaction types. It was assumed that the TF always activates the miRNA and always inhibits the target gene. This assumption could be made since loops in which the TF does not activate the miRNA and/or inhibit the target gene, will not be selected as top scoring pairs by the k-TSPs algorithm, as described below. Finally, we chose TF-miRNA and TF-target interactions which were present in at least one of the databases and miRNA-target interactions which were present in at least two databases. This resulted in 985 gene pairs which were used for predicting BLCA progression.

##### The NOTCH-MYC signaling mechanism

We used the Molecular Signature Database (MsigDB)(Liberzon et al., 2011) to retrieve gene sets associated with the regulation of the NOTCH signaling pathway or including genes up and downregulated by NOTCH. The NOTCH mechanism was constructed by pairing the genes involved in the positive regulation of NOTCH signaling pathway or genes up-regulated by NOTCH with those involved in the downregulation of the NOTCH signaling pathway or those down-regulated by NOTCH. Similarly, the MYC mechanism was constructed by pairing the genes up-regulated with those down-regulated by MYC. Finally, both mechanisms were combined into a single mechanism consisting of 78672 pairs which was further used in the TNBC classification case.

##### The cell-cell adhesion, activation, and oxygen response mechanism

We used three gene ontology (GO) biological processes to build a mechanism that can capture the biology of metastasis. These processes included: *GOBP_CELL_CELL_ADHESION, GOBP_CELL_ACTIVATION*, and *GOBP_CELLULAR_RESPONSE_TO_OXYGEN.* Unique genes were paired together resulting in 409965 gene pairs which we used as biological constraints for training classifiers to predict PCa metastatic progression.

##### Non-cancer related mechanisms

As negative controls, we designed three non-cancer related mechanisms and assessed their performance at predicting each of the three main phenotypes. First, we built a mechanism for Alzheimer disease using two gene sets including genes up- and down-regulated in the brain endothelial cells of patients with Alzheimer disease (Wu et al., 2005). Up- and down-regulated genes were paired together to form a mechanism consisting of 266 pairs. Second, we built a diabetes mechanism consisting of 6776 pairs by pairing up- and down-regulated genes in the peripheral blood monocytes from patients with diabetes at the time of the diagnosis versus 1-4 months later (Kaizer et al., 2007). Finally, we designed a mechanism consisting of 6384 pairs representing the changes in the gene expression profiles of immune cells following viral infections (Akl et al., 2007, p. 1; Dorn et al., 2005; Marshall et al., 2005). It is important to note that these three mechanisms were chosen randomly with the aim of using them as negative controls to test our hypothesis.

#### Data splitting for training and testing

In each classification case, we implemented two data splitting designs: bootstrap and cross-study validation (Xu and Goodacre, 2018) (see Figure 1). In the bootstrap design, all datasets were combined based on the set of reproducible genes. The data was then divided into 75% training and 25% testing using balanced stratification. This was done to ensure a balanced representation of the parent datasets together with important clinical and pathological variables. In the BLCA case, the clinical variables used in the stratification included age, sex, tumor grade, recurrence status, and intra-vesical therapy while in the breast cancer case, we included age, tumor grade, T and N stages. Finally, in the PCa case, we focused on age, Gleason score, tumor stage, and prostate-specific antigen (PSA) levels. Subsequently, models were trained to predict the phenotype of interest on 1000 bootstraps of the training data and their performance was evaluated on the unseen testing data using the Area Under the ROC Curve (AUC) as evaluation metric.

In the cross-study validation design, we used all but one dataset (*n-1)* for models training and the left-out dataset was used for testing. This process was repeated *n* times so that each dataset was used for testing once (see Figure 1).

#### Training and evaluating the performance of mechanistic versus agnostic models

In each classification case, we used four different algorithms: k-TSPs, RF, SVM, and XGB. Each algorithm was trained using three different model types: 1. mechanistic: using a manually curated biological mechanism in the form of pairwise comparisons (see below), 2. agnostic: using the top *2*k* differentially expressed genes (DEGs) (agnostic-genes) where k is the number of pairs used in the mechanistic models or their corresponding *k* pairwise comparisons (agnostic-pairs), 3. random genes: using a set of *2*k* randomly selected genes.

It should be noted that a pairwise comparison is based on the relative ordering of the expression of two genes. For example, in a particular sample, a given gene pair consisting of *gX* and *gY* would be assigned a value of ‘1’ if *gX* is more expressed than *gY* in that sample, and a value of ‘0’ if the opposite is true. Such pairwise comparisons were then used as features in the training process of mechanistic and agnostic-pairs models.

Importantly, all model types were trained and tested on the corresponding training and testing data, respectively. In the bootstrap approach, the AUC of each model was computed in both the training and testing data. The distribution of the AUC values of the mechanistic was plotted against those of the agnostic and random genes models to compare their average performance. In the cross-study validation, the average performance across all *n* iterations of training and testing was computed. Different metrics were used including: the AUC, accuracy, balanced accuracy, sensitivity, specificity, and Matthews correlation coefficient (MCC).

#### The k-TSPs classifier

The k-TSPs is a rank-based classification method that selects gene pairs (k) whose expression levels switch their ranking between the two classes of interest(Geman et al., 2004; Tan et al., 2005). More specifically, in the training process, if gene *X* is consistently more expressed relative to gene *Y* in samples belonging to a particular class compared to the other, it will be selected as a top scoring pair (TSP) and used for classification. In this sense, the output of this algorithm is a number of gene pairs with each voting for a specific class based on the relative ordering of expression values and the final class prediction is determined by the sum of votes. This rank assessment process used by the algorithm during the training can be applied to all genes or can be restricted to the top DEGs (agnostic) or to certain predetermined pairs chosen based on prior knowledge (mechanistic). The agnostic k-TSPs models were trained on the top DEGs by Wilcoxon rank sum test using different number of top features: the top 74, 100, 200, and 500 DEGs in the bladder and the top 25, 50, 100, 200, and 500 DEGs in both the TNBC and prostate cancer cases. The training of the mechanistic k-TSPs models was restricted to the mechanism of choice. In all cases, we restricted the number of output pairs (the final signature) to a range between 3 and 25 pairs. Finally, for each clinical problem, we obtained a matrix of binary values (*i.e.*, 0,1) representing the relative order of expression between genes pairs across individual samples (*i.e*, samples by gene pairs), which we subsequently used for training purposes with the other three prediction algorithms besides k-TSP. Such matrices of pairwise gene comparisons were generated to match the maximum number of unique starting pairs used as input to train the mechanistic k-TSPs models (37 in the bladder, 241 in the TNBC, and 50 in the prostate), selecting an appropriate number of differentially expressed genes via Wilcoxon rank sum test for the agnostic case.

#### Support Vector Machine

SVM is an algorithm that aims at identifying a hyper-plane separating data points distinctively (Noble, 2006). We trained the agnostic and mechanistic SVM models using polynomial kernel and used a repeated 10-fold cross-validation (CV) of the training data to identify the best parameter (degree, scale, and cost) values for each model. The final models were trained on the entire training data using the best parameters resulting from the repeated CV process.

#### Random Forest

RF is an ensemble ML algorithm that consists of a large number of decision trees (Breiman, 2001). Each tree in the forest votes for a specific class and the final predicted class is the one with the majority of votes. To determine the best number of variables randomly selected by the algorithm at each split (mtry), each model was tuned by the *tuneRF* function using the following parameters: *mtryStart =* 1, *ntreeTry =* 500, *stepFactor =* 1, and *improve =* 0.05. To deal with class imbalance, the final model was instructed to draw an equal number of samples from both classes for each tree. This number was set to be equal to the number of samples in the minority class of each of the training data re-samples (in the bootstrap approach) or the training data as a whole (in the cross-study validation approach).

#### Extreme gradient boosting

Similar to RF, XGB is another ensemble ML algorithm but unlike RF in which each tree is built on a random subset of predictors, XGB sub-models (sub-trees) sequentially add weight or more focus on instances with high error rates(Chen and Guestrin, 2016). We divided the training data itself into 70% “actual training” and 30% “internal validation”. We set the number of iterations to 500 with an early stopping threshold of 50 meaning that the training process will stop if the AUC in the internal validation set did not improve over 50 iterations. This step was necessary to minimize overfitting. Hyperparameters including *gamma*, *lambda*, *alpha*, and *subsample* were tuned using grid search process on the training data while stabilizing the learning rate and maximum depth. In the bootstrap analysis, this process was done on the training data before resampling. In the cross-study validation analysis, the hyperparameters were tuned on each training data partition

#### Gene Set Enrichment Analyses

To characterize the functional roles associated with the agnostic and mechanistic signatures, we performed GSEA to compute the overlap between the genes derived from the k-TSPs signatures and gene sets from the gene ontology (GO) biological processes database. In each prediction task, all unique mechanistic TSPs from the bootstrap processes were identified together with an equal number of agnostic TSPs. The GSEA analysis was performed on the genes associated with bad prognosis (positioned as gene1 in the TSPs) and those associated with good prognosis (positioned as gene2 in the TSPs), separately. P-values were calculated using Fisher’s exact test (Fisher, 1992, 1922) and were corrected using the Benjamini-Hochberg (BH) method for multiple hypotheses testing (Benjamini and Hochberg, 1995). Finally, gene sets with an adjusted p-value greater than 0.05 were considered insignificant and removed.

#### Quantification and statistical analysis

All steps of this analysis were performed using R version 4.0.3 (2020-10-10). The integrative correlation analysis was performed using the *MergeMaid* package (Cope et al., 2004). The k-TSPs models were trained using the *SwitchBox* R package (Afsari et al., 2015). The SVM models were trained using both the *Caret* (Kuhn, 2008) and *Kernlab* (Karatzoglou et al., 2004) packages. The RF and XGB models were trained using the *RandomForest* (Liaw and Wiener, 2007) and *xgboost* (Chen and Guestrin, 2016) packages, respectively. Bootstrapping (resampling with replacement) was performed using the *Boot* package (Davison and Hinkley, 1997). The values plotted in Figures 2, 4, and 6 (and supplementary figures S2:S7) represent the training and testing AUC values of the models trained on 1000 bootstraps of the training data and tested on the untouched testing data (not resampled). The values reported in Tables S2, S4, and S6 represent the average performance across all iterations of the cross-study validation process. The AUC values were computed using the prediction probabilities (RF, SVM, and XGB) or the votes (k-TSPs) returned by the classifier. The prediction probabilities were converted to predicted class labels using the optimal threshold which was determined from the training data using the ROC curve. The accuracy, balanced accuracy, sensitivity, specificity and MCC were computed by comparing the predicted class labels to the ground truth labels. The k-TSPs pairs returned from each bootstrap were ranked based on their frequency and individual genes were ranked based on their frequency in unique pairs. In the BLCA case, all mechanistic (n=93) and an equal number of agnostic pairs were used to plot the networks shown in Figure 3C-D while in the TNBC and PCa cases, the top 100 pairs were used to plot the networks shown in Figures 5 and 7C-D. All networks were built using the *igraph* R software package (Csardi and Nepusz, 2005). The size of the network vertices corresponds to twice the log2 of the gene frequency in unique pairs while the edge thickness corresponds to the log2 of frequency of the gene pair across the 1000 bootstraps. Enrichment analyses were performed using the *enrichR* package (Jawaid, 2022; Xie et al., 2021) by computing the overlap between our gene lists and gene sets from the “GO_Biological_Process_2021” database with Benjamini-Hochberg (BH) method for multiple hypotheses testing (Benjamini and Hochberg, 1995).

## Supplementary Figures

**Figure S1. An example of the coherent feed-forward loops (FFLs) used in predicting bladder cancer progression.** Here, MYC is a transcription factor repressing a downstream target gene (CD164) directly and indirectly by activating a miRNA hub (has-miR-346).

**Figure S2. The testing performance of mechanistic and agnostic models at predicting bladder cancer progression.** Models were trained on 1000 bootstraps of the training data (not shown) and evaluated on the testing data using the AUC as evaluation metric. Mechanistic models were based on the feed-forward loops mechanism (37 pairs). Agnostic models were built using either the top differentially expressed genes (top 74, 100, 200, or 500 DEGs) or the corresponding pairwise comparisons (37, 50, 100, or 250 pairs). k-TSPs: k-top scoring pairs; RF: random forest; SVM: support vector machine; XGB: extreme gradient boosting; DEGs: differentially expressed genes; AUC: Area Under the ROC Curve.

**Figure S3. Comparing the testing performance of the mechanistic and models trained on different numbers of randomly selected genes at predicting bladder cancer progression.** Models were trained on 1000 bootstraps of the training data (not shown) and evaluated on the testing data using the AUC as evaluation metric. Mechanistic models were based on the feed-forward loops mechanism (37 pairs). Random genes models were trained using different sets of randomly selected genes (74, 100, 200, and 500 genes). k-TSPs: K-top scoring pairs; RF: random forest; SVM: support vector machine; XGB: extreme gradient boosting; DEGs: differentially expressed genes; AUC: Area Under the ROC Curve.

**Figure S4. The testing performance of the mechanistic and agnostic models at predicting triple-negative breast cancer response to neoadjuvant chemotherapy.** Models were trained on 1000 bootstraps of the training data (not shown) and evaluated on the untouched testing data using the AUC as evaluation metric. Mechanistic models were based on the NOTCH-MYC mechanism (241 pairs). Agnostic models were built using either the top differentially expressed genes (top 50, 100, 200, or 500 DEGs) or the corresponding pairwise comparisons (25, 50, 100, or 250 pairs). k-TSPs: k-top scoring pairs; RF: random forest; SVM: support vector machine; XGB: extreme gradient boosting; DEGs: differentially expressed genes; TNBC: triple-negative breast cancer; NACT: neoadjuvant chemotherapy; AUC: Area Under the ROC Curve.

**Figure S5. Comparing the testing performance of the mechanistic versus random genes models at predicting triple-negative breast cancer response to neoadjuvant chemotherapy.** Models were trained on 1000 bootstraps of the training data (not shown) and evaluated on the untouched testing data using the AUC as evaluation metric. Mechanistic models were based on the NOTCH-MYC mechanism (241 pairs). Random genes models were trained using different numbers of randomly selected genes (50, 100, 200, or 500 genes). k-TSPs: K-top scoring pairs; RF: random forest; SVM: support vector machine; XGB: extreme gradient boosting; DEGs: differentially expressed genes; TNBC: triple-negative breast cancer; NACT: neoadjuvant chemotherapy; AUC: Area Under the ROC Curve.

**Figure S6. The testing performance of the mechanistic and agnostic models at predicting prostate cancer metastatic progression.** Models were trained on 1000 bootstraps of the training data (not shown) and evaluated on the untouched testing data using the AUC as evaluation metric. Mechanistic models were based on the cellular adhesion and O2 response mechanism (50 pairs). Agnostic models were built using either the top differentially expressed genes (top 50, 100, 200, or 500 DEGs) or the corresponding pairwise comparisons (25, 50, 100, or 250 pairs). k-TSPs: K-top scoring pairs; RF: random forest; SVM: support vector machine; XGB: extreme gradient boosting; DEGs: differentially expressed genes; AUC: Area Under the ROC Curve.

**Figure S7. Comparing the testing performance of the mechanistic versus random genes models at predicting prostate cancer metastatic progression.** Models were trained on 1000 bootstraps of the training data (not shown) and evaluated on the testing data using the AUC as evaluation metric. Mechanistic models were based on the cellular adhesion and O2 response mechanism (50 pairs). Random genes models were trained using different numbers of randomly selected genes (50, 100, 200, or 500 genes). k-TSPs: K-top scoring pairs; RF: random forest; SVM: support vector machine; XGB: extreme gradient boosting; DEGs: differentially expressed genes; AUC: Area Under the ROC Curve.

## Supplementary Tables

**Table S1. Gene set enrichment analyses of genes derived from the agnostic and mechanistic classifiers used to predict bladder cancer progression.** For both classifier types, GSEA was performed on all unique genes from the bootstrap design using the gene ontology biological processes. Gene1 refers to genes positioned as the 1^st^ gene in the TSPs (associated with BLCA progression) while Gene2 refers to those positioned as the 2^nd^ (associated with no-progression). Tables include only significant gene sets (adjusted p-value <0.05). Note that agnostic 1^st^ genes (gene1) were not significantly enriched in any biological processes.

**Table S2. The average performance of the agnostic and mechanistic models at predicting bladder cancer progression in the cross-study validation design.** The analysis had five iterations and in each, four studies were used for training while the fifth was used for testing. This table depicts the average training and testing performance at predicting the progression to muscle-invasive stages across the five iterations. Agnostic models were trained using either gene expression values (agnostic genes) or their pairwise comparisons (agnostic Pairs). Mechanistic models were based on the FFLs mechanism.

**Table S3. Gene set enrichment analyses of genes derived from the agnostic and mechanistic classifiers used to predict the response to neoadjuvant chemotherapy in patients with triple-negative breast cancer.** For both classifier types, GSEA was performed on all unique genes from the bootstrap design using the gene ontology biological processes. Gene1 refers to genes positioned as the 1^st^ gene in the TSPs (associated with residual disease) while Gene2 refers to those positioned as the 2^nd^ (associated with pathological complete response). Tables include only significant gene sets (adjusted p-value <0.05).

**Table S4. The average performance of the agnostic and mechanistic models at predicting the response to neoadjuvant chemotherapy in patients with triple-negative breast cancer.** Here, the analysis had seven iterations and in each, six of the seven studies were used for training while the seventh was used for testing. The table shows the average training and testing performance at predicting the response to NACT across these seven iterations. Agnostic models were trained using either gene expression values (Agnostic genes) or their pairwise comparisons (Agnostic Pairs). Mechanistic models were based on the NOTCH-MYC mechanism.

**Table S5. Gene set enrichment analyses of genes derived from the agnostic and mechanistic classifiers used to predict prostate cancer metastasis.** For both classifier types, GSEA was performed on all unique genes from the bootstrap design using the gene ontology biological processes. Gene1 refers to genes positioned as the 1^st^ gene in the TSPs (associated with metastasis) while Gene2 refers to those positioned as the 2^nd^ (associated with no-metastasis). Tables include only significant gene sets (adjusted p-value <0.05).

**Table S6. The average performance of the agnostic and mechanistic models at predicting prostate cancer metastatic progression.** The analysis included seven iterations and in each, six of the seven studies were used for training while the seventh was used for testing. The table shows the average training and testing performance at predicting metastatic events across these seven iterations. Agnostic models were trained using either individual gene expression values (Agnostic genes) or their corresponding pairwise comparisons (Agnostic Pairs). Mechanistic models were based on the cellular adhesion and O2 response mechanism.

