## Supplementary material for "Using Biological Constraints to Improve Prediction in Precision Oncology": Supp. Figures and Tables

**Supplementary Figures**

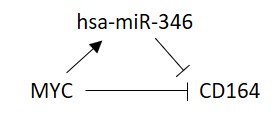

**Figure S1. An example of the coherent feed-forward loops (FFLs) used in predicting bladder cancer progression.** Here, MYC is a transcription factor repressing a downstream target gene (CD164) directly and indirectly by activating a miRNA hub (has-miR-346).

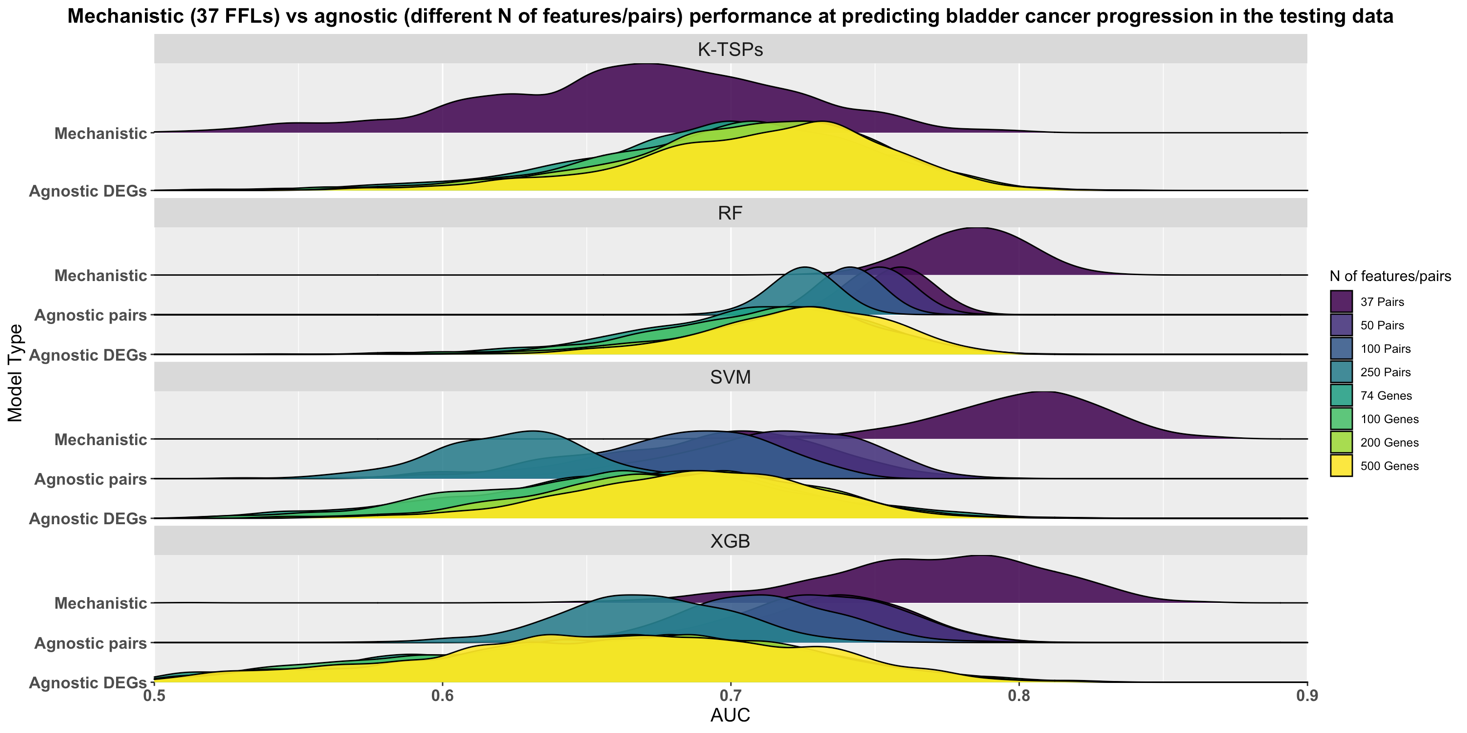

**Figure S2. The testing performance of mechanistic and agnostic models at predicting bladder cancer progression.** Models were trained on 1000 bootstraps of the training data (not shown) and evaluated on the testing data using the AUC as evaluation metric. Mechanistic models were based on the feed-forward loops mechanism (37 unique pairs). Agnostic models were built using either the top differentially expressed genes (top 74, 100, 200, or 500 DEGs) or the corresponding pairwise comparisons (37, 50, 100, or 250 pairs). k-TSPs: k-top scoring pairs; RF: random forest; SVM: support vector machine; XGB: extreme gradient boosting; DEGs: differentially expressed genes; AUC: Area Under the ROC Curve.

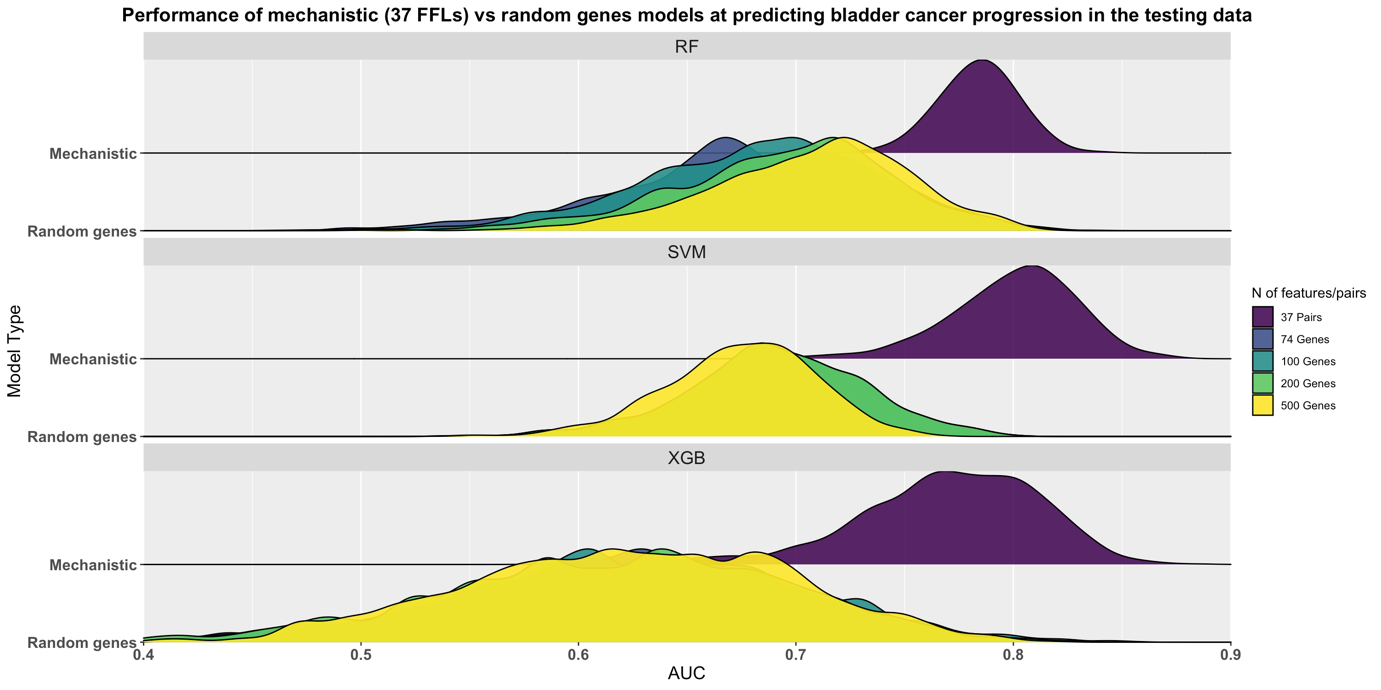
**Figure S3. Comparing the testing performance of the mechanistic and models trained on different numbers of randomly selected genes at predicting bladder cancer progression.** Models were trained on 1000 bootstraps of the training data (not shown) and evaluated on the testing data using the AUC as evaluation metric. Mechanistic models were based on the feed-forward loops mechanism (37 unique pairs). Random genes models were trained using randomly selected genes (74, 100, 200, or 500 genes). k-TSPs: K-top scoring pairs; RF: random forest; SVM: support vector machine; XGB: extreme gradient boosting; DEGs: differentially expressed genes; AUC: Area Under the ROC Curve.

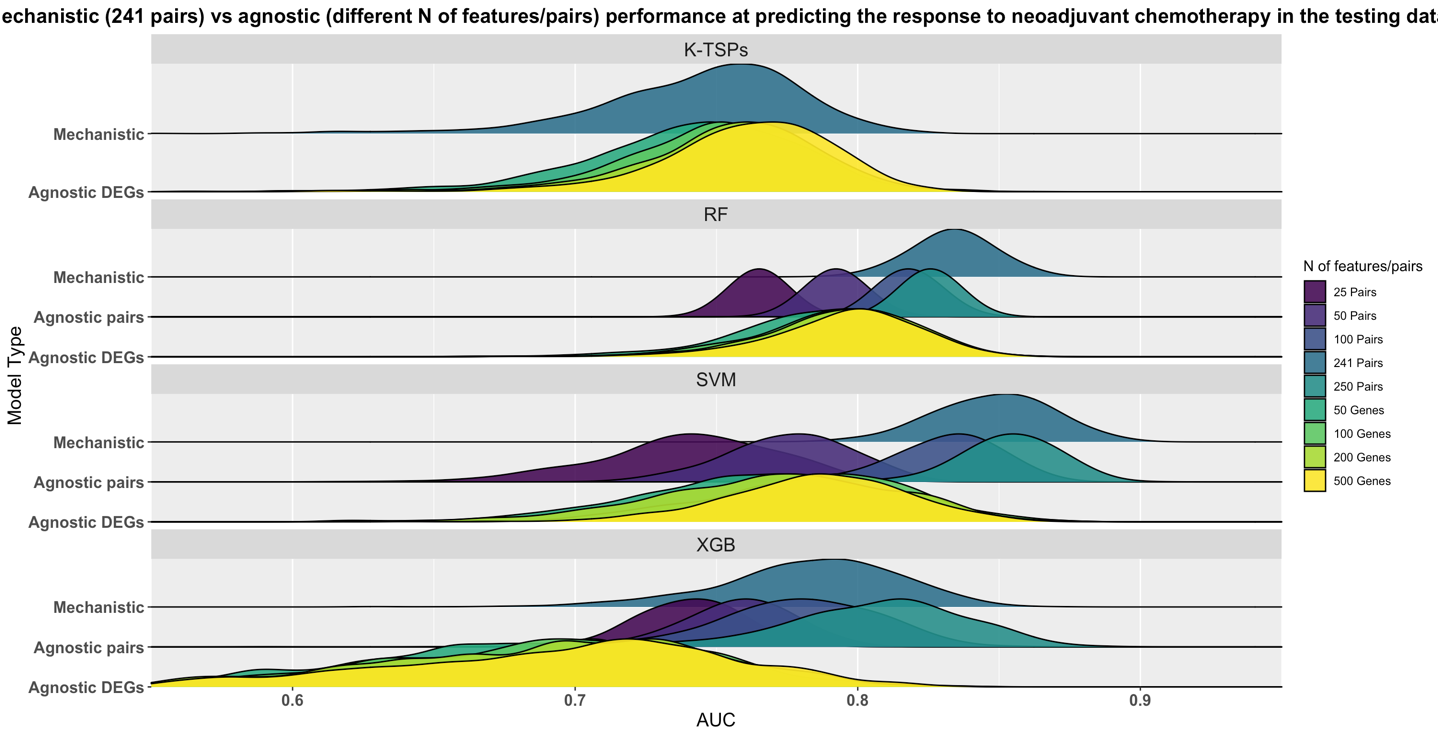

**Figure S4. The testing performance of the mechanistic and agnostic models at predicting triple-negative breast cancer response to neoadjuvant chemotherapy.** Models were trained on 1000 bootstraps of the training data (not shown) and evaluated on the testing data using the AUC as evaluation metric. Mechanistic models were based on the NOTCH-MYC mechanism (241 unique pairs). Agnostic models were built using either the top differentially expressed genes (top 50, 100, 200, or 500 DEGs) or the corresponding pairwise comparisons (25, 50, 100, or 250 pairs). k-TSPs: k-top scoring pairs; RF: random forest; SVM: support vector machine; XGB: extreme gradient boosting; DEGs: differentially expressed genes; TNBC: triple-negative breast cancer; NACT: neoadjuvant chemotherapy; AUC: Area Under the ROC Curve.

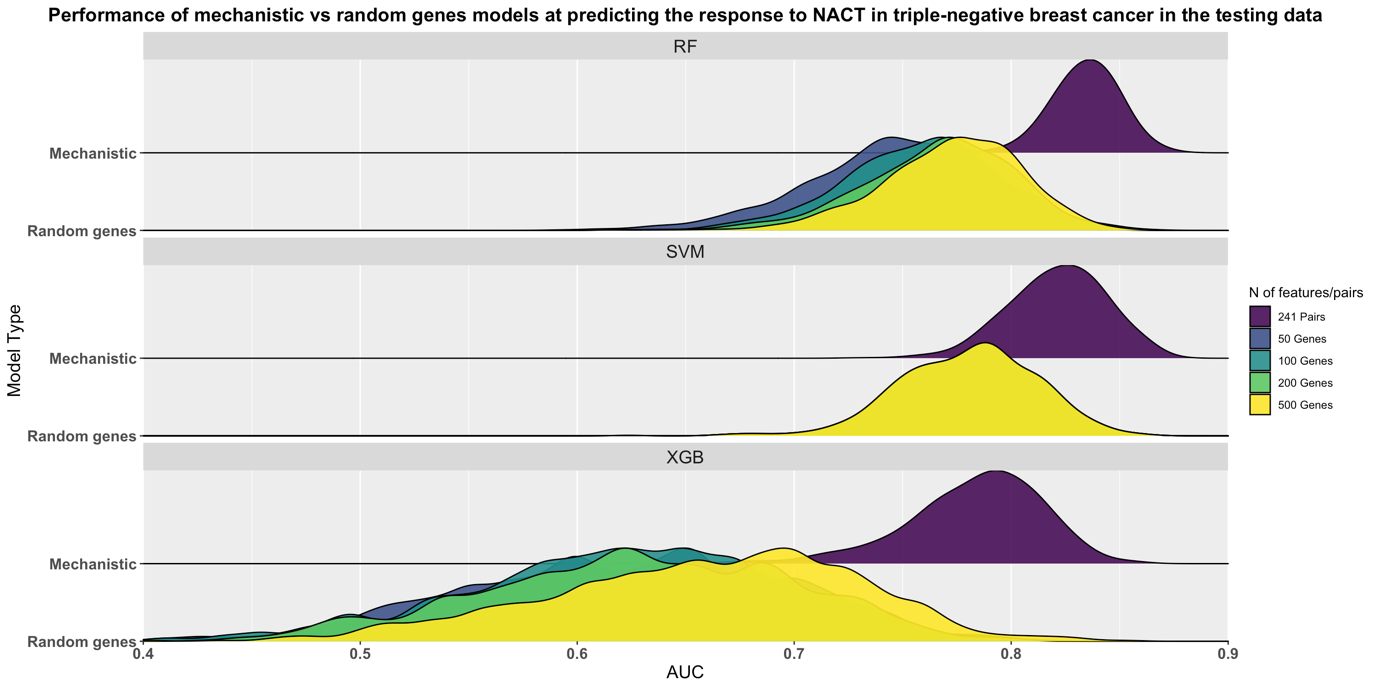

**Figure S5. Comparing the testing performance of the mechanistic versus random genes models at predicting triple-negative breast cancer response to neoadjuvant chemotherapy.** Models were trained on 1000 bootstraps of the training data (not shown) and evaluated on the untouched testing data using the AUC as evaluation metric. Mechanistic models were based on the NOTCH-MYC mechanism (241 unique pairs). Random genes models were trained using different numbers of randomly selected genes (50, 100, 200, or 500 genes). k-TSPs: K-top scoring pairs; RF: random forest; SVM: support vector machine; XGB: extreme gradient boosting; DEGs: differentially expressed genes; TNBC: triple-negative breast cancer; NACT: neoadjuvant chemotherapy; AUC: Area Under the ROC Curve.

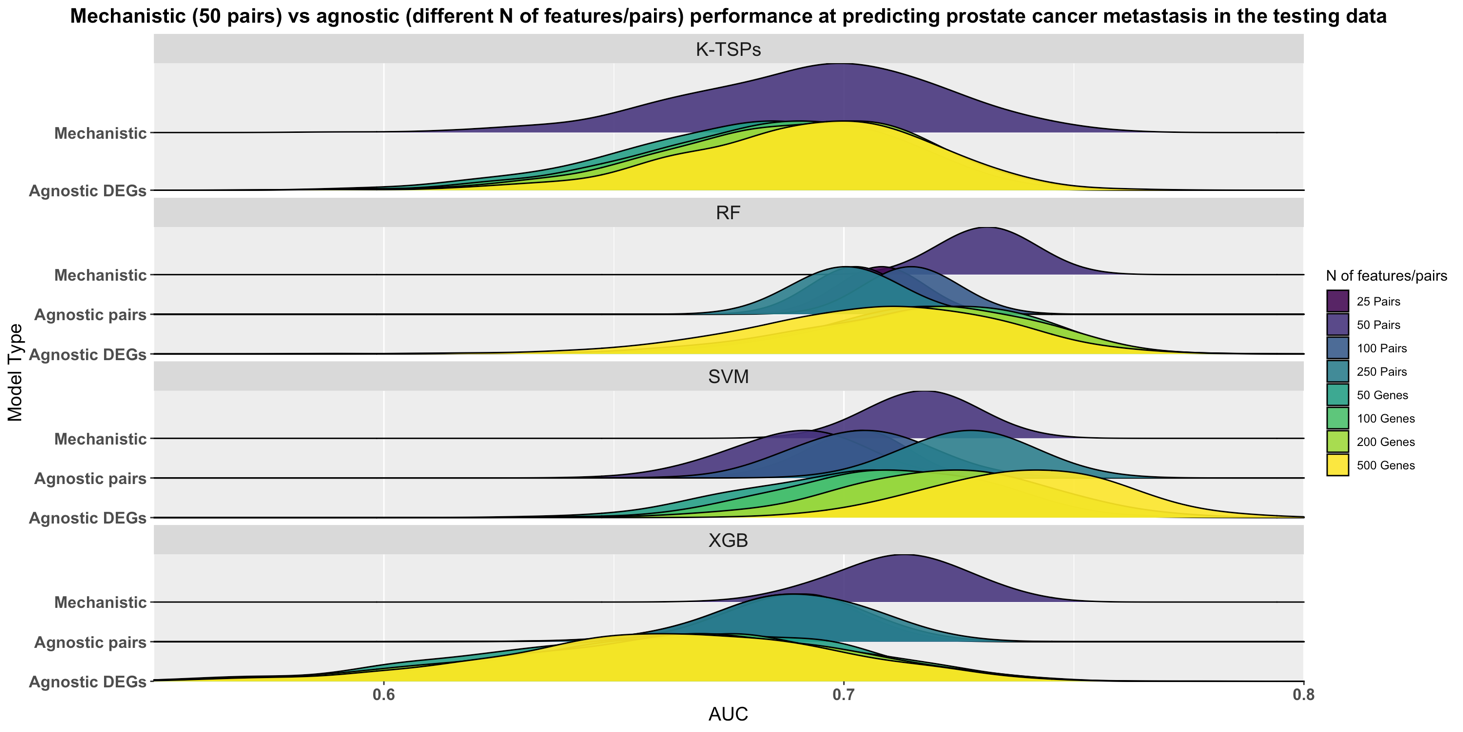

**Figure S6. The testing performance of the mechanistic and agnostic models at predicting prostate cancer metastatic progression.** Models were trained on 1000 bootstraps of the training data (not shown) and evaluated on the untouched testing data using the AUC as evaluation metric. Mechanistic models were based on the cellular adhesion and O_2_ response mechanism (50 pairs). Agnostic models were built using either the top differentially expressed genes (top 50, 100, 200, or 500 DEGs) or the corresponding pairwise comparisons (25, 50, 100, or 250 pairs). k-TSPs: K-top scoring pairs; RF: random forest; SVM: support vector machine; XGB: extreme gradient boosting; DEGs: differentially expressed genes; AUC: Area Under the ROC Curve.

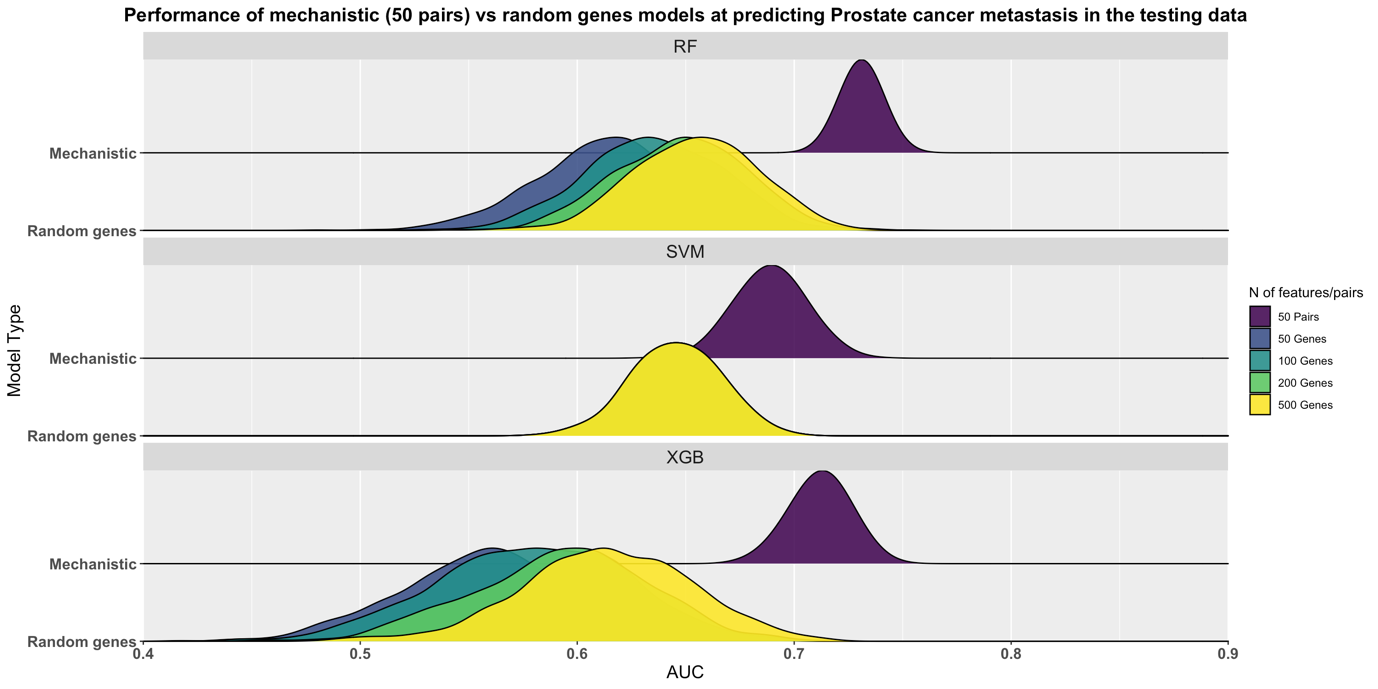
 **Figure S7. Comparing the testing performance of the mechanistic versus random genes models at predicting prostate cancer metastatic progression.** Models were trained on 1000 bootstraps of the training data (not shown) and evaluated on the testing data using the AUC as evaluation metric. Mechanistic models were based on the cellular adhesion and O_2_ response mechanism (50 pairs). Random genes models were trained using different numbers of randomly selected genes (50, 100, 200, or 500 genes). k-TSPs: K-top scoring pairs; RF: random forest; SVM: support vector machine; XGB: extreme gradient boosting; DEGs: differentially expressed genes; AUC: Area Under the ROC Curve.

**Supplementary Tables 2,4, and 6**

**Table S2. The average performance of the agnostic and mechanistic models at predicting bladder cancer progression in the cross-study validation design.** The analysis had five iterations and in each, four studies were used for training while the fifth was used for testing. This table depicts the average training and testing performance at predicting the progression to muscle-invasive stages across the five iterations. Agnostic models were trained using either gene expression values (agnostic genes) or their pairwise comparisons (agnostic Pairs). Mechanistic models were based on the FFLs mechanism.

| **Features** | **Metric** | **KTSP** ^a^ | | | **RF** | | | **SVM** | | | **XGB** | |
| --- | --- | --- | --- | --- | --- | --- | --- | --- | --- | --- | --- | --- |
|  |  | training | testing | training | | testing | raining | | testing | training | | testing |
| **Agnostic genes** | AUC  Accuracy  Bal. Accuracy  Sensitivity | NA  NA  NA  NA | NA  NA  NA  NA | 1.00 1.00 1.00  1.00 | | **0.72 0.56 0.52**  **0.18** | 1.00 1.00 1.00  1.00 | | **0.64 0.60 0.50**  **0.06** | 0.92 0.90 0.92  0.94 | | **0.64 0.64 0.55**  **0.23** |
|  | Specificity | NA | NA | 1.00 | | **0.87** | 1.00 | | **0.94** | 0.90 | | **0.86** |
|  | MCC | NA | NA | 0.99 | | **0.05** | 1.00 | | **0.00** | 0.81 | | **0.09** |
| **Agnostic pairs** | AUC  Accuracy  Bal. Accuracy  Sensitivity | 0.92 0.84 0.85  0.86 | **0.71 0.66 0.59**  **0.34** | 1.00 0.94 0.96  1.00 | | **0.71 0.60 0.55**  **0.27** | 0.95 0.89 0.90  0.91 | | **0.64 0.65 0.57**  **0.30** | 0.96 0.88 0.92  0.97 | | **0.71 0.64 0.60**  **0.38** |
|  | Specificity | 0.84 | **0.85** | 0.93 | | **0.83** | 0.89 | | **0.84** | 0.86 | | **0.82** |
|  | MCC | 0.61 | **0.19** | 0.85 | | **0.10** | 0.73 | | **0.13** | 0.73 | | **0.20** |
| **Mechanistic pairs** | AUC  Accuracy  Bal. Accuracy  Sensitivity | 0.75 0.68 0.72  0.77 | **0.69 0.57 0.62**  **0.56** | 1.00 0.95 0.97  1.00 | | **0.73 0.62 0.59**  **0.41** | 0.92 0.87 0.87  0.86 | | **0.71 0.63 0.64**  **0.53** | 0.91 0.84 0.86  0.90 | | **0.76 0.70 0.70**  **0.60** |
|  | Specificity | 0.66 | **0.69** | 0.94 | | **0.77** | 0.87 | | **0.74** | 0.83 | | **0.81** |
|  | MCC | 0.35 | **0.22** | 0.86 | | **0.14** | 0.65 | | **0.25** | 0.63 | | **0.35** |

^a^ _Note that for the K-TSPs algorithm, only pairs can be used for classification._

_K-TSPs: K-Top Scoring Pairs; RF: Random Forest; SVM: Support Vector Machine; XGB: Extreme Gradient Boosting; AUC: Area Under the ROC Curve; MCC: Matthews Correlation Coefficient._

| **Features** | **Metric** | **KTSP** ^a^ | | | **RF** | | | **SVM** | | | **XGB** | |
| --- | --- | --- | --- | --- | --- | --- | --- | --- | --- | --- | --- | --- |
|  |  | training | testing | training | | testing | training | | testing | training | | testing |
| **Agnostic genes** | AUC  Accuracy  Bal. Accuracy  Sensitivity | NA  NA  NA  NA | NA  NA  NA  NA | 1.00 1.00 1.00  1.00 | | **0.75 0.75 0.65**  **0.95** | 1.00 1.00 1.00  1.00 | | **0.79 0.76 0.69**  **0.84** | 1.00 0.99 0.99  0.99 | | **0.70 0.60 0.62**  **0.44** |
|  | Specificity | NA | NA | 1.00 | | **0.34** | 1.00 | | **0.54** | 0.99 | | **0.80** |
|  | MCC | NA | NA | 0.99 | | **0.32** | 1.00 | | **0.47** | 0.98 | | **0.25** |
| **Agnostic pairs** | AUC  Accuracy  Bal. Accuracy  Sensitivity | 0.87 0.80 0.80  0.81 | **0.70 0.63 0.62**  **0.53** | 1.00 0.99 0.99  0.98 | | **0.74 0.69 0.66**  **0.65** | 1.00 0.99 0.99  0.99 | | **0.74 0.71 0.65**  **0.72** | 0.97 0.94 0.94  0.95 | | **0.74 0.69 0.69**  **0.69** |
|  | Specificity | 0.79 | **0.70** | 1.00 | | **0.68** | 0.99 | | **0.59** | 0.93 | | **0.68** |
|  | MCC | 0.55 | **0.27** | 0.97 | | **0.36** | 0.98 | | **0.34** | 0.87 | | **0.39** |
| **Mechanistic pairs** | AUC  Accuracy  Bal. Accuracy  Sensitivity | 0.82 0.75 0.75  0.75 | **0.67 0.62 0.61**  **0.54** | 1.00 1.00 1.00  1.00 | | **0.72 0.71 0.67**  **0.74** | 1.00 0.99 0.99  0.99 | | **0.74 0.72 0.66**  **0.80** | 0.96 0.92 0.92  0.92 | | **0.71 0.68 0.69**  **0.65** |
|  | Specificity | 0.74 | **0.68** | 1.00 | | **0.60** | 0.99 | | **0.51** | 0.92 | | **0.73** |
|  | MCC | 0.48 | **0.23** | 1.00 | | **0.35** | 0.99 | | **0.33** | 0.82 | | **0.37** |

**Table S4. The average performance of the agnostic and mechanistic models at predicting the response to neoadjuvant chemotherapy in patients with triple-negative breast cancer.** Here, the analysis had seven iterations and in each, six of the seven studies were used for training while the seventh was used for testing. The table shows the average training and testing performance at predicting the response to NACT across these seven iterations. Agnostic models were trained using either gene expression values (Agnostic genes) or their pairwise comparisons (Agnostic Pairs). Mechanistic models were based on the NOTCH-MYC mechanism.

^a^ _Note that for the K-TSPs algorithm, only pairs can be used for classification._

_K-TSPs: K-Top Scoring Pairs; RF: Random Forest; SVM: Support Vector Machine; XGB: Extreme Gradient Boosting; AUC: Area Under the ROC Curve; MCC: Matthews Correlation Coefficient._

| **Features** | **Metric** | **KTSP** ^a^ | | | **RF** | | | **SVM** | | | **XGB** | |
| --- | --- | --- | --- | --- | --- | --- | --- | --- | --- | --- | --- | --- |
|  |  | training | testing | training | | testing | training | | testing | training | | testing |
| **Agnostic genes** | AUC  Accuracy  Bal. Accuracy  Sensitivity | NA  NA  NA  NA | NA  NA  NA  NA | 1.00 1.00 1.00  1.00 | | **0.77 0.73 0.60**  **0.27** | 1.00 1.00 1.00  1.00 | | **0.76 0.70 0.61**  **0.40** | 1.00 1.00 1.00  1.00 | | **0.73 0.70 0.64**  **0.41** |
|  | Specificity | NA | NA | 1.00 | | **0.94** | 1.00 | | **0.83** | 1.00 | | **0.87** |
|  | MCC | NA | NA | 1.00 | | **0.27** | 1.00 | | **0.24** | 1.00 | | **0.26** |
| **Agnostic pairs** | AUC  Accuracy  Bal. Accuracy  Sensitivity | 0.80 0.71 0.72  0.75 | **0.75 0.67 0.70**  **0.65** | 1.00 0.99 0.99  1.00 | | **0.77 0.71 0.68**  **0.50** | 0.84 0.76 0.77  0.81 | | **0.76 0.65 0.68**  **0.65** | 0.84 0.76 0.76  0.78 | | **0.76 0.67 0.68**  **0.60** |
|  | Specificity | 0.69 | **0.74** | 0.98 | | **0.85** | 0.73 | | **0.71** | 0.75 | | **0.76** |
|  | MCC | 0.41 | **0.34** | 0.97 | | **0.33** | 0.51 | | **0.31** | 0.50 | | **0.32** |
| **Mechanistic pairs** | AUC  Accuracy  Bal. Accuracy  Sensitivity | 0.78 0.70 0.71  0.75 | **0.75 0.65 0.69**  **0.63** | 1.00 0.99 0.99  1.00 | | **0.76 0.71 0.67**  **0.48** | 0.88 0.80 0.81  0.84 | | **0.74 0.69 0.67**  **0.60** | 0.79 0.71 0.73  0.77 | | **0.76 0.66 0.68**  **0.62** |
|  | Specificity | 0.68 | **0.74** | 0.98 | | **0.86** | 0.78 | | **0.74** | 0.69 | | **0.73** |
|  | MCC | 0.40 | **0.32** | 0.97 | | **0.33** | 0.59 | | **0.29** | 0.43 | | **0.30** |

**Table S6. The average performance of the agnostic and mechanistic models at predicting prostate cancer metastatic progression.** The analysis included seven iterations and in each, six of the seven studies were used for training while the seventh was used for testing. The table shows the average training and testing performance at predicting metastatic events across these seven iterations. Agnostic models were trained using either individual gene expression values (Agnostic genes) or their corresponding pairwise comparisons (Agnostic Pairs). Mechanistic models were based on the cellular adhesion and O_2_ response mechanism.

^a^ _Note that for the K-TSPs algorithm, only pairs can be used for classification._

_K-TSPs: K-Top Scoring Pairs; RF: Random Forest; SVM: Support Vector Machine; XGB: Extreme Gradient Boosting; AUC: Area Under the ROC Curve; MCC: Matthews Correlation Coefficient._
